# Genomic signatures associated with transitions to viviparity in Cyprinodontiformes

**DOI:** 10.1101/2022.05.25.493474

**Authors:** Leeban H. Yusuf, Yolitzi Saldívar Lemus, Peter Thorpe, Constantino Macías Garcia, Michael G. Ritchie

**Affiliations:** Centre for Biological Diversity, School of Biology, University of St Andrews, St Andrews, UK; School of Medicine, Mackenzie Institute for Early Diagnosis, University of St Andrews, North Haugh, St Andrews, UK; Instituto de Ecologia, Universidad Nacional Autónoma de México, Ciudad Universitaria, Circuito exterior s/n anexo al Jardín Botánico C. P. 04510, Mexico City D. F, Mexico

**Keywords:** convergent evolution, molecular evolution, comparative genomics, viviparity

## Abstract

The transition from oviparity to viviparity has occurred independently over a hundred times across vertebrates, presenting a compelling case of phenotypic convergence. However, whether repeated, independent evolution of viviparity is driven by redeployment of similar genetic mechanisms and whether these leave a common genetic signature in genomic divergence remains unknown. Whilst investigations into the evolution of viviparity have demonstrated striking similarity among the genes and pathways involved across vertebrate groups, quantitative tests for genome-wide convergence provide ambivalent answers. Here, we investigate molecular convergence during independent transitions to viviparity across an order of ray-finned freshwater fish (*Cyprinodontiformes*). We assembled *de novo* and publicly-available genomes of viviparous and oviparous species to quantify molecular convergence across coding and non-coding regions. We found no evidence for an excess of molecular convergence in amino acid substitutions and rates of sequence divergence, implying independent genetic changes are associated with these transitions. However, statistical power and biological confounds (hemiplasy and introgression) could constrain our ability to detect correlated evolution. We therefore also identified candidate genes with potential signatures of molecular convergence in viviparous *Cyprinodontiformes* lineages. While we detected no evidence of positive or relaxed selection for these genes in branches associated with the evolution of viviparity in *Cyprinodontiformes*, motif-enrichment and gene ontology analyses suggest transcriptional changes associated with early morphogenesis, brain development and immunity occurred alongside the evolution of viviparity. Overall, our findings indicate that an excess of molecular convergence, at any level, is not strongly associated with independent transitions to viviparity in these fish.

## Introduction

The extent of genomic homology during the convergent evolution of complex phenotypes is a major question in evolutionary genetics, but poorly resolved. While the independent acquisition of similar traits in lineages that do not share recent common ancestry is eloquent evidence for parallel adaptive evolution, such phenotypic convergence may involve similar genetic changes (Castoe *et al*., 2009; Martin and Orgogozo, 2013; Stern, 2013; Mu *et al*., 2015), but sometimes distinct molecular and developmental processes are involved (Steiner *et al*., 2008; Berens, Hunt and Toth, 2014; Natarajan *et al*., 2016; Murugesan *et al*., 2022). Recent attempts to understand the genetic bases of phenotypic convergence have demonstrated the importance of comparative phylogenomic approaches to identify genetic changes responsible for independent trait transitions across different taxonomic groups (Smith *et al*., 2020). Even when common genetic factors are involved, they may be diverse, ranging from parallel amino acid changes (Johanson *et al*., 2000; Sugawara *et al*., 2005; Kuittinen *et al*., 2008), gene family evolution (Christin *et al*., 2007), convergent sequence divergence in protein-coding genes (Chikina, Robinson and Clark, 2016), and convergent sequence divergence in regulatory regions (Sackton *et al*., 2019). However, disentangling confounding factors from convergent evolution has proven difficult (Parker *et al*., 2013; Thomas and Hahn, 2015; Zou and Zhang, 2015). The challenge is often exacerbated in cases of complex phenotypic traits, where convincing signals of molecular convergence may be especially difficult to detect (Corbett-Detig *et al*., 2020; Yusuf *et al*., 2020), and previous comparative genomic analyses that have searched for convergence have been limited as they either (a) focus only on convergent amino acid changes, (b) are restricted to convergent shifts in gene-wide evolutionary rate, or (c) lack empirical null models to determine whether neutral processes may explain the observed signals of molecular convergence. Few attempts to characterize molecular convergence have jointly addressed the potential role of phylogenetic incongruence and hemiplasy in confounding patterns of molecular convergence (Sackton *et al*., 2019; Corbett-Detig *et al*., 2020; Hibbins, Gibson and Hahn, 2020).

The convergent evolution of viviparity provides a tractable system to explore complex, independent trait transitions and the consistency of molecular adaptations. Viviparity has independently evolved from oviparity (egg laying) more than 150 times across diverse vertebrate groups, with fish and squamate reptiles accounting for most transitions (Blackburn, 1999). Relatively recent reversals back to oviparity and variable modes of reproduction within species have been documented, suggesting transitions in parity may be relatively labile (Blackburn, 2015; Recknagel, Kamenos and Elmer, 2018; Recknagel and Elmer, 2019; Recknagel *et al*., 2021). The evolution of viviparity involves a suite of complex morphological, physiological and developmental changes including egg retention, internal fertilization, immunotolerance, internal embryonic development and nutrient and gas exchange (Van Dyke, Brandley and Thompson, 2014). These multiple adaptations are in part explained by the maternal-foetal interface and degree of parity which differs considerably amongst viviparous species, with embryos displaying either some degree of maternal dependency (matrotrophy) or relative metabolic autonomy (lecithotrophy) (Blackburn, 1992).

Previous analyses of genomics in relation to viviparity have largely focused on genomic and transcriptomic transitions linked to mammalian viviparity, despite it only containing a single transition in reproductive mode (Lynch and Wagner, 2008; Lynch *et al*., 2008; Emera *et al*., 2012; Kin *et al*., 2016; Nnamani *et al*., 2016). Recent genomic analyses in seahorses, pipefish and lizards which have repeatedly evolved viviparity have identified modules of genes that show similarity in homology and function likely to be involved in transitions to viviparity in mammals (Whittington *et al*., 2015; Gao *et al*., 2019; Roth *et al*., 2020). In particular, shifts in the expression of genes involved in hormonal regulation, tissue remodelling and nutrient exchange, as well as convergent loss of genes involved in adaptive immune response were found to coincide with the evolution of viviparity in these taxa. However, despite increasing availability of genomic data covering independent transitions to viviparity, there have been few explicit tests of genome-wide molecular convergence linked to viviparity and associated adaptations (Gao et al. 2019; Roth et al. 2020; van Kruistum et al. 2021; Recknagel et al. 2021). Recently, Foster et al. (2022) analysed the transcriptomes of the placenta of eight vertebrates (not including fish) and concluded that there was no significant overlap in gene involvement, so each placenta had evolved from unique changes in gene expression networks.

*Cyprinodontiformes* is an order of small, freshwater ray-finned fish that show considerable diversity in reproductive mode and associated adaptations (Wourms and Lombardi, 1992). Within the order, the evolution of viviparity has been linked to bursts of diversification explained by changes in associated life-history traits, sexual selection and sexual conflict (Helmstetter *et al*., 2016). In poecilids, phylogenetic comparative methods have demonstrated that the evolution of matrotrophy and placentation is associated with sexual conflict and sexual selection (Pollux *et al*., 2014; Furness *et al*., 2019). Similarly, comparative genomic analyses within the order have specifically focused on understanding the genetic context of placental evolution, uncovering parallels with mammalian viviparity and between independent acquisitions of placentae in poecilids (Guernsey et al. 2020; van Kruistum, et al. 2021). The *Goodeidae*, a family of mostly viviparous ray-finned fish within the order *Cyprindontiformes* have received less attention. The *Goodeinae* are proposed to have become viviparous after diverging from oviparous species of the family in the early Miocene, before subsequent diversification of matrotrophic adaptations (Webb *et al*., 2004). However, the role of sexual selection and conflict as a driver of viviparity and matrotrophy in Goodeinae remains unclear (Ritchie *et al*., 2005; Saldivar Lemus *et al*., 2017).

Here, we utilize *de novo* genome assemblies and publicly available genomes to identify genomic features associated with the evolution of viviparity. Specifically, we use a phylogenomic comparative framework consisting of poecilids from four different genera (*Poecilia*, *Gambusia*, *Poeciliopsis* and *Xiphophorus*) and goodeids from two subfamilies (*Goodeinae* and the *Empetrichthyinae*), alongside oviparous pupfish (*Cyprinodon* and *Orestias*) and mummichog (*Fundulus*) genomes. Our phylogenetic framework is thought to contain two independent transitions from oviparity to viviparity (Helmstetter *et al*., 2016). By comparing 16 viviparous and 5 oviparous genomes and leveraging their phylogenetic context, we sought to identify genomic regions associated with the convergent evolution of viviparity in Cyprinodontiformes at three levels by assessing: (a) convergent changes in amino acids, (b) evolutionary rate change across protein-coding genes and (c) evolutionary rate change in conserved, non-coding regions. Additionally, to test whether genome-wide convergent amino acid changes and evolutionary rate divergence could be explained by neutral processes, we computed empirical null models by randomizing and retesting foreground branches in our phylogenetic framework. As with some previous analyses looking for quantitative evidence of convergent molecular changes (Thomas and Hahn, 2015; Zou and Zhang, 2015; Gao *et al*., 2019; Corbett-Detig *et al*., 2020), we find no evidence of an excess of genome-wide convergence in protein-coding genes that may explain transitions to viviparity. We do identify genetic variation associated with shifts to viviparity and discuss how detecting signals of convergence may be confused by incomplete lineage sorting.

## Results

### Genome assembly and annotation

Genomes assembled in this project, including raw reads and annotation are available from NCBI using the accession codes in Supplementary Table 1. Metrics for *de novo* genome assemblies are summarized in Supplementary Table 2.

### Reconstructing the molecular phylogeny of *Cyprindontiformes*

The species tree generated (Supplementary Figure 1) has an almost identical topology to the pruned tree obtained from Rabosky et al. (2018) (Figure 1) with two exceptions. First, our tree inferred *Girardinichthys multiradiatus* as being more distantly related to *Goodea atripinnis* and *Xenoophorus captivus*, as opposed to *Xenotaenia resolanae* as seen in Rabosky et al (2018). Secondly, *Poecilia formosa* and *Poecilia latipinna* form a species pair in the species tree inferred in this study, which is inconsistent with both Rabosky et al. (2018) used in the analysis, and Warren et al. (2018). In both cases of phylogenetic disagreement, branches in our species tree show high levels of gene discordance (33.13% and 48.19%, respectively)(Supplementary Figure 1). However, these ambiguous branches are both background branches so are unlikely to impact our analyses. We used the pruned tree from Rabosky et al (2018) for all subsequent analyses (Figure 1).

**Figure 1:**
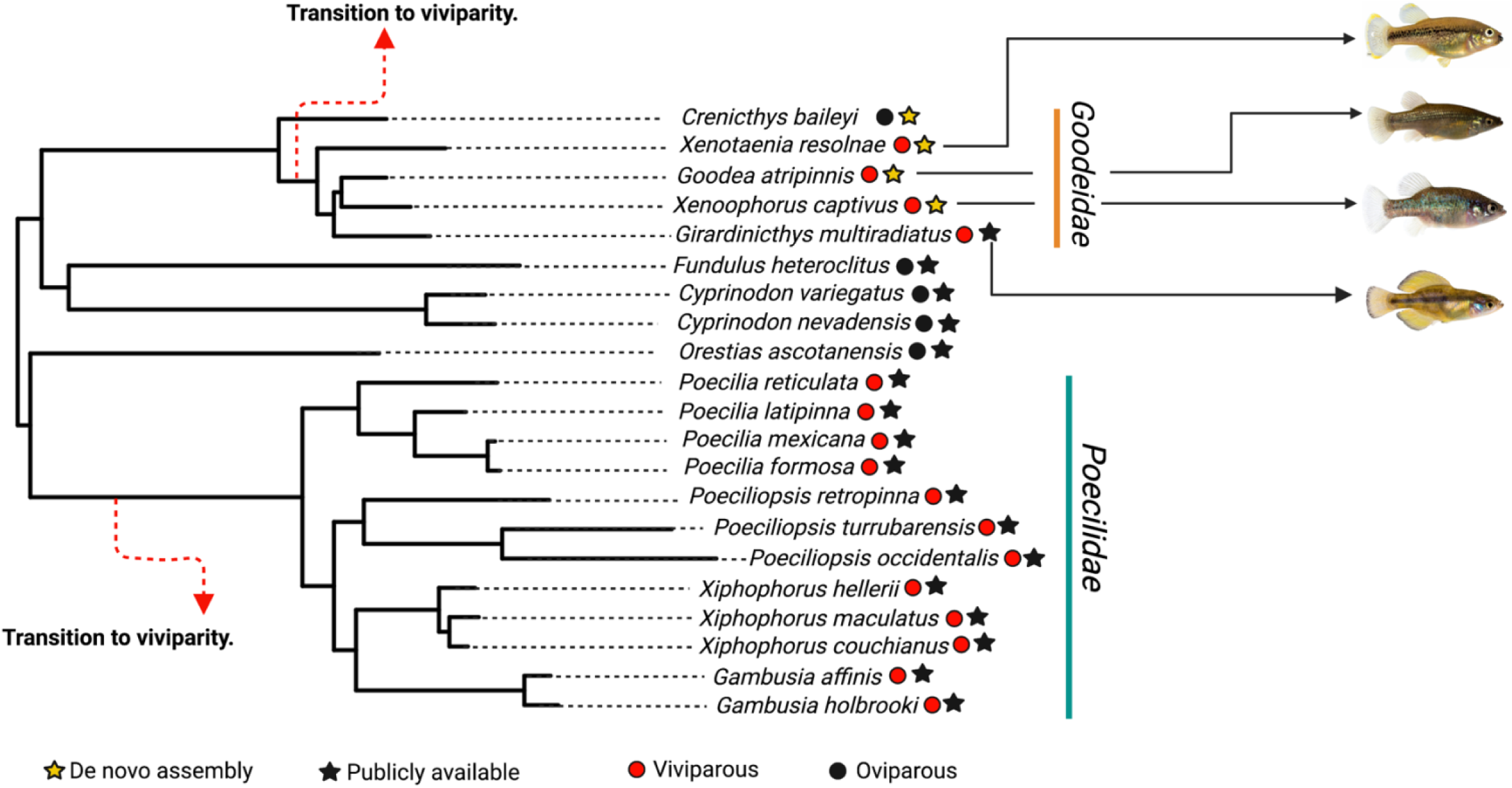
Pruned phylogenetic tree of the order *Cyprinodontiformes* from Rabosky et al (2018). This is the phylogeny used in comparative analysis of molecular convergence. Both hypothesized transitions to viviparity used as foreground branches in the analysis are shown, alongside character states (either viviparous or oviparous) for each species denoted by circles near species names. Stars denote whether genomes were assembled here or whether they were publicly available.

### No significant evidence for genome-wide convergent evolution at the amino acid level

In total, 17,572 orthologous protein alignments were generated and filtered, all containing sequence data for each of the 21 species. We found that 2,038 convergent amino acid changes occurred in foreground branches where viviparity is inferred to have evolved. However, these are fewer than those expected under empirical null models. In both empirical null models, foreground branches were randomized along the tree, so that these represented branches that were phenotypically and phylogenetically distinct. The number of convergent amino acid changes observed were 9,018 and 13,937 in the first and in the second empirical null models respectively. Hence, our data show no evidence for an excess in genome-wide convergent evolution on foreground branches associated with shifts to viviparity. This relates exclusively to the quantity, not to the “quality” of the changes, and whilst we found that signals of apparent convergence at single amino-acid sites are phylogenetically widespread and not restricted to foreground branches, a subset of these genes may nevertheless be linked to the emergence of viviparity. To explore this, we examined the functions of genes showing evidence of convergent amino acid changes in foreground branches, by performing gene ontology overrepresentation analysis for biological processes against human and zebrafish backgrounds. Using the human background dataset, we found an enrichment of genes in important signalling pathways known to be involved in embryogenesis, namely Wnt signalling (fold enrichment=3.68; FDR = 3.25e-02) and Hippo signalling (fold enrichment=12.49 ; FDR = 3.79e-02) (see reviews; Steinhart & Angers 2018; Davis & Tapon 2019) (Supplementary Figure 2a). Using the zebrafish background, we found an enrichment of terms involved in embryogenesis, for example, embryonic viscerocranium morphogenesis (fold enrichment = 4.03; FDR = 4.97e-02) and embryonic organ morphogenesis (fold enrichment = 2.34; FDR = 4.56e-02) (Supplementary Figure 2b). We also found evidence of convergent amino acid evolution (FDR < 0.05) in estrogen receptor 1 (GPER1), prolactin (PRL), and luteinizing hormone/choriogonadotropin receptor (LHCGR), suggesting rewiring of hormonal regulation associated with the evolution of viviparity. Additionally, we found evidence of a convergent amino acid change in CD74, a gene critically important in the MHC II pathway and for the recognition of non-self peptides. In both pipefish and seahorses (Roth *et al*., 2020), convergent exon loss and divergence of exons in CD74 were associated with the evolution of male pregnancy. Further functional studies are required to fully characterize the role of convergent evolution in the structure and function of adaptive immune responses in viviparous Cyprinodontiformes.

As well as searching for convergent amino acid changes, we also assessed relative and correlated rates of protein evolution with phylogenetic switches to viviparity, i. e. proteins that changed in evolutionary rate alongside viviparity. We found that 532 genes showed a correlation (p < 0.05) between relative rate of protein evolution and the evolution of viviparity, though none were significant after correcting for multiple testing. Of these 532 genes, 310 showed slower evolutionary rates at foreground compared to background branches, and 222 faster evolutionary rates. However, when compared to empirical null distributions we again failed to find a significant excess of genes with correlated sequence divergence in our experimental group compared to the controls (ANOVA: df=3, F=1.8027, p = 0.2862).

In order to examine the functions of these 532 genes further, we surveyed mean expression across the placentae of 14 mammalian species and found that 254 genes (with annotations) showed non-negligible expression with mean fragments per kilobase per million reads (FPKM) > 1 (χ^2^ = 1.0827, df = 1, p-value = 0.2981). Additionally, those that show accelerated or conserved relative evolutionary rates in branches where viviparity evolved also show higher than average mammalian placental expression compared to both a control gene set (W = 163083, p-value = 0.0076) and to the background mammalian placenta expression data (W = 3117536, p-value < 2.2e-16) (Figure 2a). Genes showing appreciable expression in placental mammals and correlated sequence divergence include fibroblast growth factors, tyrosine-protein kinase (JAK1), myogenic factor 6 (MYF6), fibronectin (FN1), integrin beta-1 (ITGB1), insulin receptor substrate 1 (IRS1) and notably, SMAD2, a downstream protein in the transforming growth factor β signalling family (Figure 2b and Supplementary Figure 2). In Mammals, members of transforming growth factor β signalling family have been shown to be involved in ovulation (Lee *et al*., 2007), decidualization (Lee *et al*., 2007; Fullerton *et al*., 2017), implantation (Clementi *et al*., 2013; Peng, Monsivais, *et al*., 2015) and placentation (Peng, Fullerton, *et al*., 2015).

**Figure 2:**
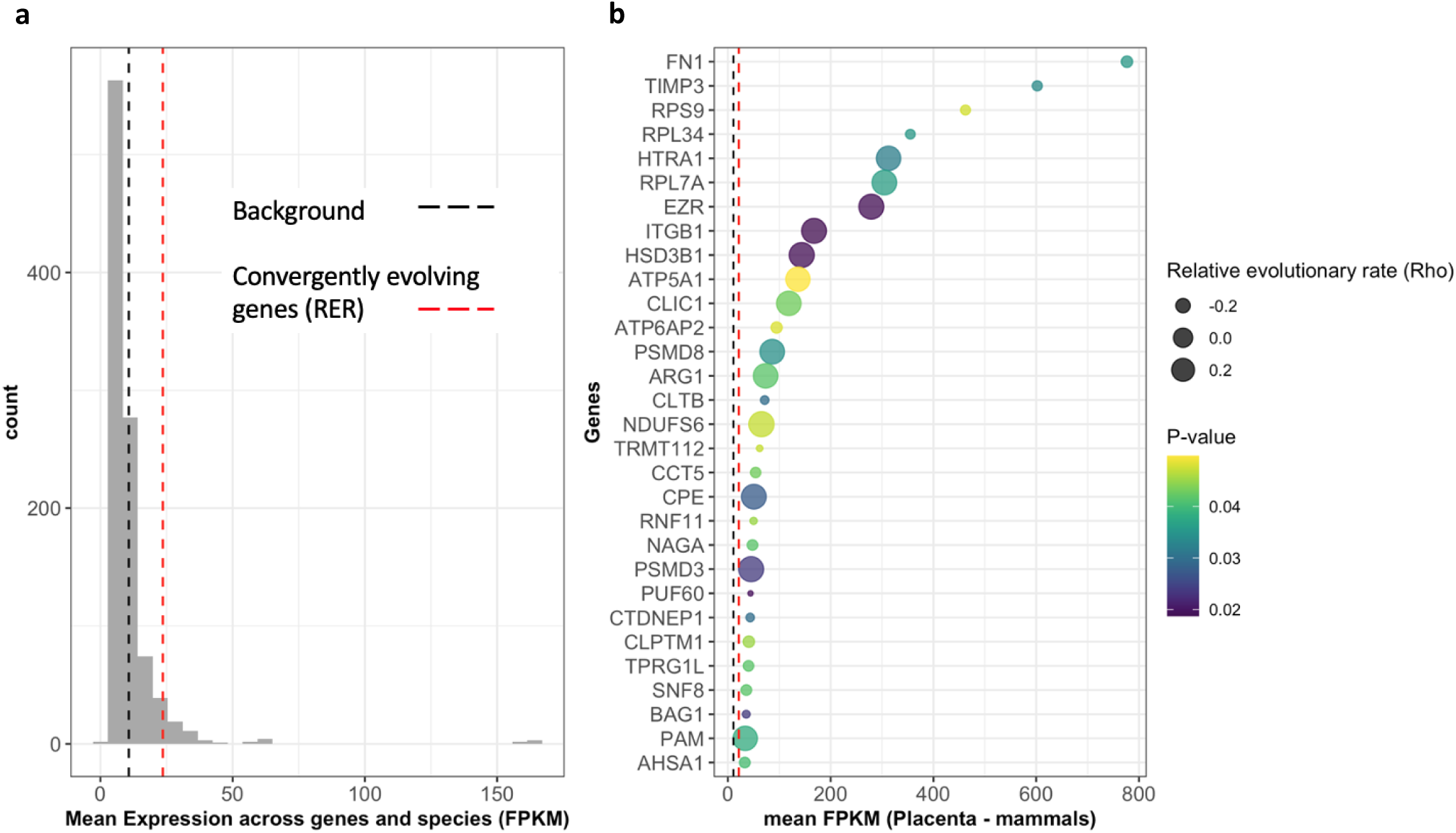
Genes with correlated sequence divergence in branches where transition to viviparity occurred. **a)** Empirical null distribution of control gene set, with black dashed line showing mean mammalian placental expression across all genes and red dashed line showing mean expression for convergently evolving genes**. b)** The top 30 genes with correlated sequence divergence ordered by mean mammalian placental expression (FPKM). Unadjusted p-value indicates significance of relative evolutionary rate.

By overlapping the results from the two previous analyses, we found 59 genes that showed evidence of both (1) a convergent amino acid change at foreground branches, and (2) a significant association between relative evolutionary rate and the trait change to viviparity on foreground branches(Figure 3a). To determine whether positive selection has acted on any of these, we conducted gene-wide and branch-site tests using BUSTED and aBSREL (Murrell *et al*., 2015; Smith *et al*., 2015). Neither test revealed evidence of positive selection in any of the genes at branches involving switches to viviparity. However, in total, 48 of the 59 genes show evidence of positive selection on at least one internal or terminal branch leading to a viviparous lineage and in at least one site. To confirm this, aBSREL was used to detect evidence of positive selection on branches leading to viviparous lineages (Figure 3b). This identified the same 48 (out of the 59) genes as showing evidence of positive selection (P < 0.05). Finally, after multiple testing correction, we found no genes that showed evidence of relaxed selection and only two genes — Delta and Notch-like epidermal growth factor-related receptor (DNER) and dedicator of cytokinesis 2 (DOCK2) —showed an intensification of selection in branches where viviparity evolved in Cyprinodontiformes (Table 1).

**Figure 3:**
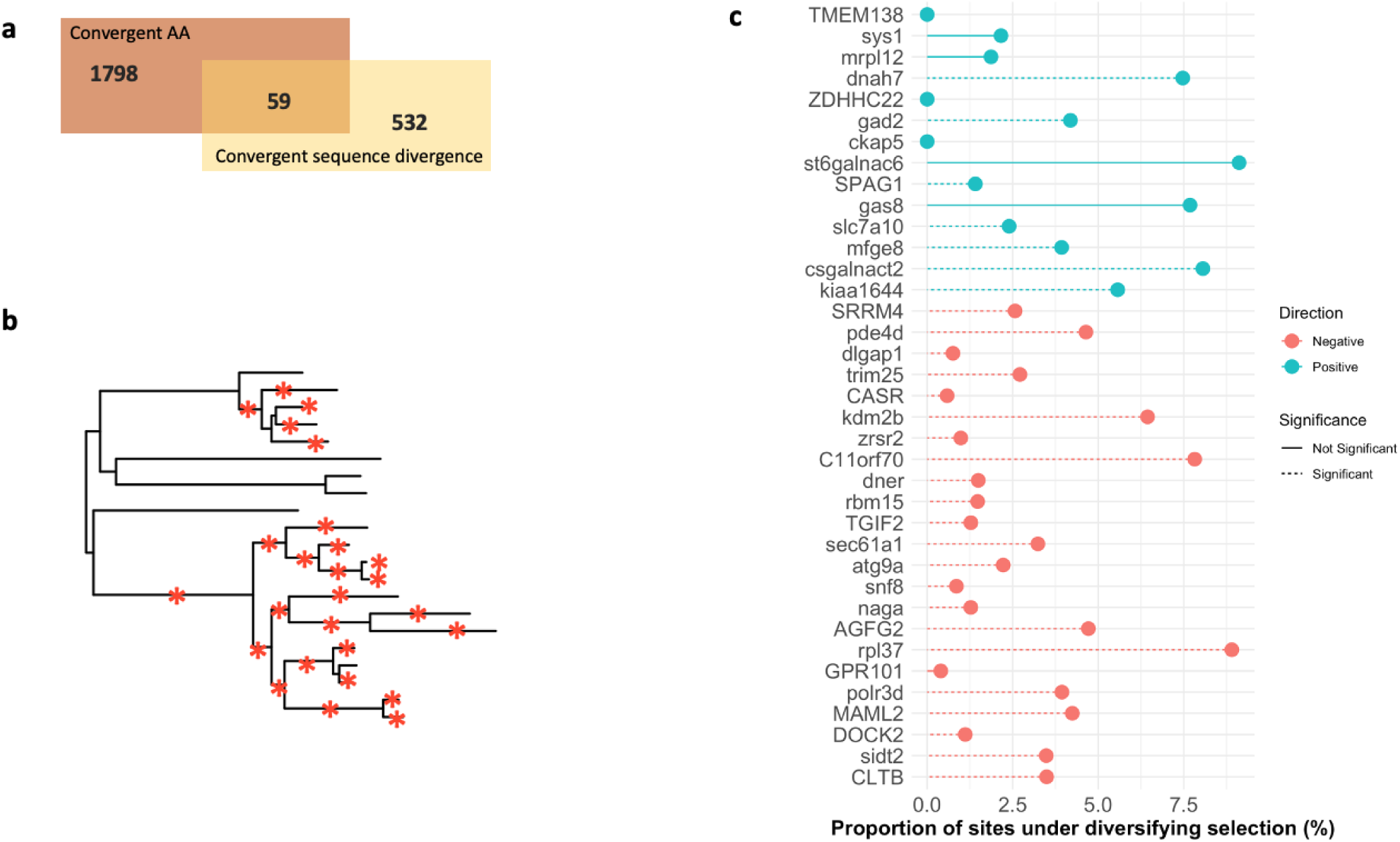
**a)** Overlap between genes with convergent amino acid substitutions and convergent evolutionary rates. **b)** The phylogenetic tree of Cyprinodontiformes marked with red asterisks denoting branches that were individually tested for positive selection. **c)** Estimates of proportion of sites under diversifying selection for genes showing both a convergent amino acid substitutions and convergent evolutionary rate. Genes with dashed lines are those with evidence of positive selection in at least one branch. Genes categorized by whether they show constrained or accelerated rates compared to background branches in RER analysis.

**Table 1:** Nineteen genes showing evidence of convergent molecular evolution, positive selection and which are expressed in mammalian placentas and in the maternal follicles of two Poeciliopsis species.

| Gene name | P-value (BUSTED) | P-value (RELAX) | Found in mammalian placentas | Found in maternal follicle of *Poeciliopsis* species | Evidence for intensification or relaxation of selection? |
| --- | --- | --- | --- | --- | --- |
| CSGALNACT2 | 0.0029 | 0.0672 | Yes | Yes | No intensification or relaxation |
| DOCK2 | <0.0001 | <0.0001 | Yes | Yes | Intensification |
| ATG9A | <0.0001 | 0.302 | Yes | Yes | No intensification or relaxation |
| MAML2 | <0.0001 | 0.6401 | Yes | Yes | No intensification or relaxation |
| DNER | 0.0001 | 0.0004 | Yes | Yes | Intensification |
| SIDT2 | <0.0001 | 0.4686 | Yes | Yes | No intensification or relaxation |
| RBM15 | <0.0001 | 0.8877 | Yes | Yes | No intensification or relaxation |
| RPL37 | 0.0019 | 0.2578 | Yes | Yes | No intensification or relaxation |
| KDM2B | <0.0001 | 1 | Yes | Yes | No intensification or relaxation |
| CKAP5 | 0.0217 | 0.0323 | Yes | Yes | Relaxation |
| DNAH7 | <0.0001 | 0.281 | Yes | Yes | No intensification or relaxation |
| POLR3D | 0.0001 | 0.019 | Yes | Yes | Intensification |
| PDE4D | <0.0001 | 0.1176 | Yes | Yes | No intensification or relaxation |
| TRIM25 | 0.001 | 0.3251 | Yes | Yes | No intensification or relaxation |
| CLTB | <0.0001 | 0.0187 | Yes | Yes | Intensification |
| SPAG1 | <0.0001 | 0.6066 | Yes | Yes | No intensification or relaxation |
| SLC7A10 | <0.0001 | 0.6595 | Yes | Yes | No intensification or relaxation |
| SNF8 | <0.0001 | 0.1004 | Yes | Yes | No intensification or relaxation |
| NAGA | 0.0027 | 0.012 | Yes | Yes | Relaxation |

Despite showing some evidence of both positive selection and convergent molecular evolution, the 59 genes found here may be evolving due to selective pressures for traits that are correlated with the evolution of viviparity (e.g. attributes that, while not responsible for any “viviparity trait”, may nevertheless facilitate the evolution of viviparity). To consider this we examined whether genes amongst those 59 were expressed in relevant tissues. Expression data of maternal follicles of two species within the *Poeciliopsis* clade (*P. retropinna* and *P. turrubarensis*) are available (Guernsey et al. (2020) and also for placental tissue of 14 mammal species (Armstrong et al. (2017). In mammalian placental tissue, 31 of the 59 genes showed non-negligible expression (FPKM >1) (χ^2^ = 0.15254, df = 1, p-value=0.6961). (Supplementary Figure 4). In the *Poeciliopsis*, 21 genes showed non-negligible expression in either species (*P.retropinna* FPKM > 3.2877, *P.turrubarensis* FPKM > 6.1751, following cut-offs from Guernsey et al. 2020) and these genes also showed concordant non-negligible expression in mammals.

Altogether, 19 genes (32%) showed (1) a convergent amino acid change, (2) an association between relative evolutionary rate and convergent switches to viviparity, (3) evidence of positive selection on lineages where viviparity has evolved, (4) appreciable expression in either of the maternal follicles of two *Poeciliopsis* species and (5) appreciable expression in the placenta of 14 mammals (Table 1). These include dedicator of cytokinesis 2 (DOCK2), which is known to activate RAC1/RAC2 genes required for implantation (Kunisaki *et al*., 2006; Grewal *et al*., 2008), and TRIM25, a gene involved in innate immune response against viral infection, mediating oestrogen action and whose downregulation during embryogenesis in medaka has been shown to result in apoptosis (Gack *et al*., 2007; Dong *et al*., 2012; Zhang *et al*., 2015).

### Conserved non-coding elements are convergently accelerated in viviparous lineages

Alongside protein-coding genes, regulatory regions can drive convergent evolution of phenotypes and may be under less functional constraints than protein-coding genes (Carroll 2008). To identify putative cis-regulatory elements that evolved concordantly in foreground branches, we extracted conserved non-coding regions using a neutral model where substitution rates were estimated from four-fold degenerate sites, and then tested whether any conserved non-coding region showed acceleration of sequence divergence at foreground branches where the transition to viviparity occurred. We found 733,850 conserved non-coding regions evolving significantly slower than four-fold degenerate sites across the alignment. After correcting for multiple testing, only 245 of these showed significant sequence acceleration at both foreground branches (Supplementary Table 6). These showed a significant enrichment of transcription-factor binding motifs. These transcription factors were enriched in gene ontology terms related to embryonic organ morphogenesis and embryo development ending in birth or egg hatching (Supplementary Table 7).

To determine what genes may be regulated by the 245 accelerated non-coding elements, we extracted the closest neighboring genes to each accelerated non-coding element. We found 181 genes neighboring at least one accelerated non-coding element, with 32 genes showing more than one accelerated non-coding element nearby. Genes showing a higher density of accelerated non-coding elements nearby include RNF144a, an E3 ubiquitin ligase and FGFR4, a fibroblast growth factor receptor. To examine potential functions of all 181 genes, we performed gene ontology analysis of biological and molecular terms and found a significant enrichment of terms related to synapse development, development of the sympathetic nervous system, neuron development and telencephalon development (Figure 4b). Additionally, we found significant enrichments for transcription factors and genes with both E-box (enhancer box) domains and HMG-box domains (Figure 4a). These genes include GATA3, SREBF2, NEUROG1, TCF4, POU3F3 and PRXX1. Altogether, motif-enrichment and gene ontology analyses suggest transcriptional changes associated with brain development evolved alongside the evolution of viviparity.

**Figure 4:**
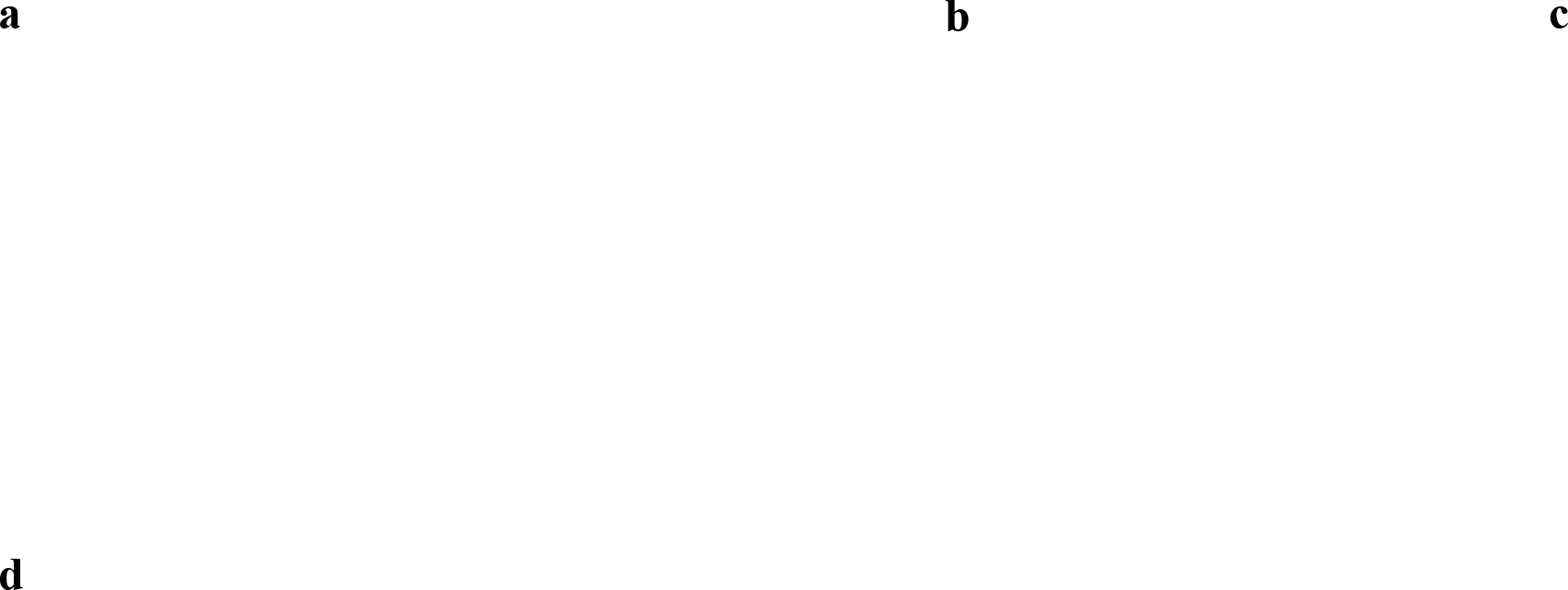
Evidence of convergent evolution in accelerated, conserved, non-coding elements. **a**) Protein domains of transcription factors associated with putative binding sites in 245 accelerated non-coding elements**. b)** Gene ontology (biological terms) for genes nearby accelerated non-coding elements. **c)** Number of accelerated conserved, non-coding elements nearby each gene. Genes with more than five elements nearby are labelled. **d)** Sequence divergence in foreground branches in conserved, non-coding elements shown across scaffolds arranged by linkage group (based on guppy genome). Non-coding elements in red show significant (FDR<0.05) acceleration in branches where the transition to viviparity occurred.

### Exploring the potential influence of incomplete lineage sorting

To understand the potential effect of hemiplasy (due to incomplete lineage sorting alone), we estimated the likely number of genetic changes associated with viviparity by simulating 10^8^ loci along the inferred *Cyprinodontiformes* species tree. Using HeIST, we converted branch lengths (in substituitions per site) to coalescent units using gene concordance estimates and the regression approach detailed in Hibbins et al. (2020). We found that for focal cases – where simulated nucleotides match character traits in the species tree – a scenario where 1) all viviparous taxa are grouped together in a single monophyletic clade, and where 2) a transition to viviparity is explained by a single mutation, is more common (41/79 of focal loci) than homoplasy (38/79 of focal loci). These results suggest that under the assumption that transitions to viviparity are underpinned by a simple genomic architechture, there is an almost equal probability that viviparity in *Cyprinodontiformes* may be explained by a single transition (ancestral polymorphism) rather than two.

## Discussion

Comparative studies of vertebrate viviparity, stimulated by increasing genomic resources, development of phylogenetic comparative methods and growing evidence of molecular convergence in general (Smith *et al*., 2020), are beginning to ask if there is a common genetic underpinning of viviparity (Lynch et al 2008; Kin et al 2016). Some have shown remarkable convergent evolution of genes and pathways involved in independent transitions to viviparity across both closely-related and distantly-related species, however other studies are suggest that species can evolve striking parallel adaptation by redeployment of different genes, including the vertebrate placenta (Foster et al. 2020). Here, we have considered the role of molecular convergence in the repeated evolution of viviparity in *Cyprinodontiformes*. We found no evidence for an excess of genome-wide molecular convergence in amino acid changes or in rates of protein evolution in explaining the convergent evolution of viviparity. Our results suggest convergent evolution on protein-coding genes is not strongly associated with viviparity in *Cyprindontiformes*, and may also involve independent genetic changes even within this group. . However, we succeeded in identifying a small subset of candidate genes that show evidence of convergent evolution and are homologous to genes essential in mammalian viviparity, suggesting a small number of genes may be required for the evolution of viviparity, but perhaps too few to result in a strong signal of parallel evolution overall .

### The evolution of viviparity is not associated with a significant excess of genome-wide molecular convergence

An important source of support for invoking convergent evolution in comparative genomic analysis is finding convergent amino acid substitutions or convergent sequence divergence in independent lineages that are greater than expected by chance (Zhang, 2003; Storz, 2016). However, the detection of genome-wide molecular convergence has proved contentious. For example, comparative surveys identifying genome-wide convergence in mammalian lineages that independently evolved echolocation were shown to have fewer parallel substitutions than comparisons between echolocating and non-echolocating mammals (Parker *et al*., 2013; Thomas and Hahn, 2015; Zou and Zhang, 2015). Similarly, behavioural and morphological phenotypic convergence in anole lizards is not associated to parallel substitutions or rates of amino acid change, suggesting either (a) difficulty in detecting genome-wide convergence, or (b) that the same loci are not repeatedly recruited (Corbett-Detig *et al*., 2020). Similarly, in a comparison of oviparous and viviparous lizards, evidence for convergent amino acid change was minimal, with most consistent differences observed in gene expression (Gao et al. 2019). Independent acquisitions of placentotrophy in *Poecilidae* showed a preponderance of protein-coding genes undergoing shifts in sequence divergence in placental species compared to non-placental species (van Kruistum, et al. 2021). In our study, we did not find a higher incidence of convergence at any level examined – in fact we found a higher incidence of molecular convergence in our null models.

However, a small subset of genes in each of our analyses show striking resemblance to genes critically important for mammalian viviparity, and they belong to gene families previously implicated in viviparity-related adaptations. For example, we found evidence of convergent amino acid substitution in prolactin. In Eutherian (placental) mammals, prolactin is produced throughout pregnancy, stimulates cell proliferation and differentiation, and maintains production of progesterone and relaxin (Soares, Konno and Alam, 2007). Additionally, prolactin expression was previously observed in the maternal follicle of matrotrophic species of *Poeciliopsis*, but not in lecithotrophic species of *Poeciliopsis*, suggesting a potential role for prolactin in the evolution of matrotrophy in *Poecilidae* and *Goodeinae* (Guernsey *et al*., 2020). We also found evidence of convergent evolution in genes critical to adaptive immunity that are also associated with the evolution of viviparity. Changes to adaptive immunity in viviparous lineages may be responsible for maternal immunotolerance towards developing embryos. In viviparous seahorses and pipefish, there is similar loss of genes related to the MHC II pathway and rapid sequence evolution of CD74, an invariant chain of MHC II involved in preventing premature binding (Roth *et al*., 2020). Here, we found evidence of convergent substitution in a gene involved in the MHC II pathway in viviparous lineages, though it is unknown whether this substitution is neutral or under selection.

### No evidence of positive selection on branches associated with the transition to viviparity

We assessed sequence divergence associated with viviparity and compared signatures of convergence with placental expression data across mammals. We found that a number of genes showing concordant sequence divergence show elevated expression in all mammalian placentas, suggesting that genes implicated in mammalian pregnancy may be repeatedly recruited in non-mammalian viviparity. These include HTRA1, EZR, ITGB1 and HSD3B1 which have roles in foetal adhesion, foetal growth, invasion of endometrial stromal cells and modulation of progesterone (Ornek *et al*., 2008; Shimodaira *et al*., 2012; Burkin *et al*., 2013; Nishimura *et al*., 2014). Some genes may be housekeeping ones with ubiquitous expression that are involved in general cellular maintenance and metabolism. For example, we identify a number of ribosomal proteins (RPS9, RPL34, RPL7A) that are highly expressed across mammalian placentas and show signals of concordant sequence divergence.

We found 59 genes with evidence of both a convergent amino acid change and correlated sequence divergence. Whilst some of these show evidence of positive selection in at least one branch in the phylogeny, none showed evidence of positive selection in both branches associated with the transition to viviparity. These results mirror patterns of convergent evolution of placentae in *Poecilidae* (van Kruistum, et al. 2021). Similarly, an analysis of marine mammals found that very few genes showing concordant sequence divergence also showed evidence of positive selection (Chikina, Robinson and Clark, 2016). Instead, genes showing accelerated sequence divergence in marine mammals were probably subject to relaxed constraints (Chikina, Robinson and Clark, 2016). Here, we found no evidence of relaxed selection in the 59 genes on branches associated with the transition to viviparity. Since all 59 genes were tested for positive selection in branches associated with the transition to viviparity, as well as all internal and terminal branches leading to viviparous species, correcting for multiple testing may have raised the bar to detect potential signals of positive selection only on branches associated with the transition to viviparity.

### Can biological confounds explain independent transitions to viviparity in Cyprinodontiformes?

As well as statistical power, another potential confound in detecting parallel evolution arises due to collateral evolution. Collateral evolution describes the evolution of phenotypic convergence via genetic variation that was either (a) the result of some introgression event or (b) occurred via the involvement of an ancestral polymorphism and incomplete lineage sorting (Stern, 2013). Here, we addressed the potential for collateral evolution to explain patterns of molecular convergence. Surprisingly, when simulating hemiplasy under ILS alone, we found an almost equal probability of viviparity in *Goodeinae* and *Poecilidae* having a single genetic basis. However, there are a number of important caveats. First, we do not include all independent transitions to viviparity, in particular, we do not include the evolution of viviparity in *Anablepidae* (Helmstetter *et al*., 2016), which likely results in underestimating the probability of a homoplastic origin in our analysis. Secondly, the approach used assumes the genetic architecture of simulated traits are monogenic, but a simple genetic architecture is unlikely given the necessary and complex adaptations required for viviparity to evolve. Thirdly, whilst we simulated a large number of loci, only 79 focal loci matched the species tree and were considered as either hemiplastic or homoplastic. As a result, whether hemiplasy is likely to play a major role in our ability to detect convergent evolution probably remains an open question.

### Accelerated non-coding regions associated with the evolution of viviparity show evidence of functionality

Beyond assessing molecular convergence in coding regions, we also sought to identify non-coding regions undergoing accelerated sequence divergence in branches associated with the transition to viviparity. We identified a small number of conserved non-coding regions showing accelerated sequence divergence and nearby genes that may be associated with these regions. These accelerated non-coding loci showed an enrichment of transcription factor binding motifs, suggesting functionality of identified accelerated non-coding loci and potential rewiring of gene expression associated with the evolution of viviparity. In particular, some of the genes identified were located nearby multiple accelerated non-coding loci, for example, FGFR4, a fibroblast growth factor receptor with evidence of expression in the human placenta (Anteby *et al*., 2005). We also detected 149 genes with only a single accelerated non-coding loci nearby with known roles in mammalian viviparity (Supplementary Table 8). For example, GATA3 has previously been identified as a critical and conserved component of placental development in mammals (Home *et al*., 2017; Gerri *et al*., 2020). However, when genes were considered together via gene ontology analysis, we found an enrichment for genes with a potential role in mediating the development of the central nervous system. Whilst these analyses do not clearly implicate particular pathways, they suggest a cohort of genes with potentially broad functions which may have undergone changes in gene expression during the evolution of viviparity in Goodeids and Poecilids.

## Conclusions

The study of convergent evolution of genes underlying traits such as complex morphologies often highlights contrasts between consistent changes in cohorts of consistent genes versus redeployment of independent gene networks. Our analyses indicate that this might be a somewhat artificial distinction. We find both a lack of a strong quantitative signal of concerted parallel changes in the evolution of viviparity in the *Cyprinodontiformes*, but convincing evidence of the involvement of a few genes important to the evolution of viviparity across these and additional vertebrates. More independent transitions to viviparity will help resolve statistical issues in detecting consistent gene changes, but out study suggests more attention to potential biological confounds such as hemiplasy and introgression is also needed.

## Methods and Materials

### Sample collection

Muscle tissue samples were obtained from adult males of *Goodea atripinnis, Xenotaenia resolanae*, *Xenophoorus captivus* and *Crenichthys baileyi*. *G. atripinnis, X. resolanae* and *X. captivus* individuals were descendants of fish captured in Michoacán, Jalisco and San Luis Potosí states (Mexico) under SEMARNAT permit SGPA/DGVS/00824/20. A muscle tissue sample of a male of the oviparous goodeid *Crenichthys bailey*i was obtained from Kees de Jong.

### Whole-Genome Sequencing

Extracted DNA was sent to Novogene (Beijing, China), for library preparation and sequencing. Sequencing libraries were generated using the NEBNext® DNA Library Prep Kit (New England Biolabs, USA) following the manufacturer’s instructions. The genomic DNA was sheared to a size of 350bp, then the fragments were end-polished, A-tailed, and ligated with the NEBNext adapter (New England Biolabs™, USA) for Illumina sequencing. Resulting libraries were analysed for size distribution with 2100 Bioanalyzer (Agilent™) and quantified using real-time PCR. Paired-end sequencing was performed on an Illumina NovaSeq 6000 system (Illumina™ Inc.) using the v1.0 reagents for sequencing.

### Genome assembly

Raw reads were interrogated for quality using FastQC (Andrews, 2010) then quality trimmed using Trimmomatic (version 0.38) (Q15) (Bolger, Lohse and Usadel, 2014). The quality controlled reads were then assembled using spades (version 3.14.1) (Prjibelski *et al*., 2020). Blobtools version 1.0 (Laetsch and Blaxter, 2017) was used to identify and remove contaminant contigs, the assembly was compared to the non-redundant database (Genbank nt) using BLASTn (megablast). Contigs identified as fungal, bacterial, plantal or viral were removed, yielding a contamination free final unpolished assembly. The assemblies were then scaffolded using 1 iteration SSPACE (v3.0) (Boetzer *et al*., 2011) with BWA. Finally, 3 iterations of Pilon (version 1.23) (Walker *et al*., 2014) was performed to polish the assembly. At all stages of assembly BUSCO (Simão *et al*., 2015) version 1.1b was used to assess the relative completeness of the assembled genomes using Eukaryotic odb9 models.

Gene annotation was predicted using funannotate version 1.8.0. Soft-masked genomes were generated as described in Thorpe et al. (2018). All RNAseq separately for each species was concatenated into a single left and right read file, explicitly not mixing or mapping reads from different species. These was quality trimmed at Q30 using trimmomatic and normalised using bbnorm (Last modified October 19, 2017) (https://jgi.doe.gov/data-and-tools/bbtools/bb-tools-user-guide/bbnorm-guide/). The reads were mapped to the corresponding genome using STAR (v2.5.3a) (Dobin *et al*., 2013) with --outFilterMultimapNmax 1 --outFilterMismatchNmax 4 parameters to only return uniquely mapping reads. The resulting bam was passed to trinity (v2.8.4) (Haas *et al*., 2013) for genome guided RNAseq assembly with -- genome_guided_min_coverage 4 to remove contigs with low support. The resulting genome guided RNAseq assembly was subjected to 2 iterations of filtering with Transrate (v1.0.3) (Smith-Unna *et al*., 2016).The resulting assembly was then used by funannotate to run and train PASA (v2.4.1) (Haas *et al*., 2008) and Augustus (v3.3.3) (Stanke *et al*., 2006). The resulting gene predictions were functionally annotated using PFAM (https://pfam.xfam.org/), InterProScan (https://www.ebi.ac.uk/interpro/search/sequence/) and Eggnog (Huerta-Cepas *et al*., 2019).

### Genome alignment and filtering

Additional genomes of closely-related viviparous and oviparous *Cyprinodontiformes* species (Figure 1) were downloaded from RefSeq and GenBank (Supplementary table 1). To determine evolutionary relationships and divergence times, a time-calibrated phylogeny for *Cyprinodontiformes* was retrieved from Rabosky et al. (2018) and pruned using the ape (v5.5) package in R to retain the relevant species used here (Paradis *et al*., 2003; Paradis and Schliep, 2019). To produce a whole-genome alignment of the 21 genomes, we aligned all genomes to a reference genome (*G. multiradiatus*; Du et al. 2022) using LAST aligner (Kiełbasa *et al*., 2011; Hamada *et al*., 2017), followed by chaining and netting using scripts and utilities from the UCSC browser source code (Miller *et al*., 2003). Finally, a multiple whole-genome alignment of the 21 species was built from the reciprocal-best nets using MULTIZ (Blanchette *et al*., 2004), with *G. multiradiatus* as the reference.

Protein coding sequences were extracted from the multiple whole-genome alignment using the *G. multiradiatus* genome annotation (Du et al., 2022) and MafFilter (v1.3.0) (Dutheil, Gaillard and Stukenbrock, 2014). To obtain aligned orthologous amino acid sequences, we initially aligned coding sequences using MAFFT (v7.471) with default settings (Katoh *et al*., 2002, 2005; Katoh and Standley, 2013; Nakamura *et al*., 2018). Subsequently, MACSE v.2 was used to mask frameshifts and stop codons and the resulting nucleotide sequences converted into amino acid sequences (Ranwez *et al*., 2018). Finally, Divvier (options: partial -divvygap - mincol 21) was used to retain only clusters of high-confidence amino-acid columns in alignments with evidence of shared homology, removing variable columns in alignments that were deemed low-confidence (Ali et al. 2019). For codon sequences, MACSE v.2 was used to trim homologous fragments from the beginning and end of alignments and then MACSE v.2 was used to align sequences. Codon alignments were translated into amino acids and subsequently converted into final codon alignments using PAL2NAL, specifying the removal of incomplete codons and gaps (Suyama, Torrents and Bork, 2006).

### Phylogenetic reconstruction of species tree and estimating the probability of hemiplasy

In order to infer species relationships, we retrieved 1,044 single-copy orthologs from a BUSCO analysis of all 21 species (performed using the *Actinopterygii* gene set) (Simão *et al*., 2015). Species-specific proteomes consisting of these single-copy orthologs were then used to infer orthogroups and a species tree using Orthofinder2 and IQ-TREE2 with the following parameters: -M msa -T iqtree (Nguyen *et al*., 2015; Emms and Kelly, 2018). To quantify phylogenetic discordance, we utilized this species tree and gene trees inferred from 16,941 filtered amino acid alignments using IQ-TREE2 with the parameters: -m LG+G4 -nt 16.

We quantified the probability of hemiplasy in our phylogenetic framework using HeIST (Hibbins, Gibson and Hahn, 2020) which uses coalescent simulations to estimate the probability of hemiplasy and homoplasy under a multispecies network coalescent model, given a species tree and gene/site concordance factors. Specifically, we used the species tree inferred above with gene concordance factor estimates to convert branch lengths into smoothed coalescent units. Within HeIST, *ms* and *Seq-gen* were used to simulate 10^8^ loci along the species tree, and only focal loci reflecting the specific character states in the species tree were considered (Rambaut and Grassly, 1997; Hudson, 2002). Since mutation rates for species within the phylogeny have not been estimated, we followed the rationale of Hibbins et al. (2020) and used a vertebrate-general estimate of 0.005 per 2*N* generations.

### Protein coding genes associated with the transition to viviparity

To detect convergent amino acid changes in lineages that have evolved viviparity, we utilized the TDG09 program, which compares a null model assuming homogenous substitution patterns for each site in the alignment across all branches in the phylogeny, with a model which assumes non-homogenous substitution patterns between ‘foreground’ branches where viviparity is assumed to have evolved, and ‘background’ branches (Tamuri *et al*., 2009). All amino acid alignments that passed filtering were tested using the species tree with ‘foreground’ or ‘background’ annotations ascribed to each species in the tree (groups VI OV, for viviparity and oviparity, respectively). To determine whether an excess of genome-wide convergent amino acid changes has occurred in foreground branches, empirical null models of the phylogenetic framework were conducted. Foreground branches were randomized in two different control tests and all sites were retested. Only sites with a FDR < 0.05 (Benjamini and Hochberg, 1995) were considered to be convergently evolving.

Convergent genetic changes may involve the same changes in single amino acids, but also coding sequences may show correlated divergence across the entire protein. We used RERconverge to test the correlation between relative rates of protein evolution and convergent trait evolution at foreground branches on the species tree (Chikina, Robinson and Clark, 2016; Kowalczyk *et al*., 2019). We generated phylogenetic trees for all amino acid alignments using the LG amino acid matrix and, by fixing the tree topology based on the species tree so that only branch lengths were estimated (Le and Gascuel, 2008; Schliep, 2011), then transformed into relative evolutionary rates to determine whether, for a given branch, genes have evolved faster or slower than the background rate.

Branch-specific relative change rates were then used in a correlation analysis with binary trait classifications of either ‘viviparity’ or ‘oviparity’, where viviparous species were designated as foreground branches. As a control, we randomly sampled two foreground branches across the phylogeny and re-tested all genes to determine whether we observed an excess of molecular convergence. A weighted correlation was used to correct for heteroscedasticity. Weighted correlations were carried out using Kendall’s Tau and multiple testing correction was performed using the Benjamini-Hochberg procedure (Benjamini and Hochberg, 1995).

To test whether genes showing an association with viviparity in our fish samples were expressed in mammalian placental tissue, expression data for mammalian placental tissues across 14 species were retrieved from Armstrong et al. (2017). To assess whether genes showing significant relative evolutionary rates (SRER) in foreground branches where viviparity evolved also showed higher than expected expression in the mammalian placenta, we compared mean expression (as FPKM) between SRER genes and an empirical null distribution generated by permuting (1000 iterations) expression levels of randomly sampled genes. Additionally, expression data for *Poeciliopsis turrubarensis* (lecithotrophic) and *Poeciliopsis retropinna* (matrotrophic) were retrieved from Guernsey et al. (2020). Specifically, genes were listed as either expressed in the maternal follicles of *P. turrubarensis*, *P. retropinna*, or both.

### Testing for evidence of positive selection and shifts in selection pressure

To test for evidence of positive selection along viviparous lineages, we utilized two different codon substitution models: BUSTED and aBSREL (Murrell *et al*., 2015; Smith *et al*., 2015). Specifically, we tested for positive selection in genes that showed both at least one convergent amino acid substitution and a significantly different relative evolutionary rate in foreground branches. BUSTED tests for evidence of gene-wide positive selection – that is, whether there has been diversifying selection on at least one foreground branch and for at least one site in the alignment using the dN/dS ratio. An unconstrained model with three rate classes (which sites are assigned to) was compared with a null (constrained) model where diversifying selection is disallowed, using a likelihood ratio test. Alongside this, aBSREL was used to test if positive selection has occurred on a subset of sites in specified foreground branches, i.e. a branch-site test. For both tests, foreground branches were specified as the branches where the hypothesized transition to viviparity occurred (Figure 1). Additionally, we tested for positive selection on all internal and terminal branches leading to viviparous lineages that are not representative of the transition from oviparity to viviparity. Finally, we used RELAX (Wertheim *et al*., 2015) to test whether, at branches where the transition to viviparity occurred (2 foreground branches as observed in Figure 1), genes that show evidence of convergent evolution have experienced relaxation or intensification of selection.

To determine the putative functions of genes, we used eggNOG-mapper to annotate genes and PANTHER to perform gene ontology for biological processes using human and zebrafish reference databases (Huerta-Cepas *et al*., 2017; Mi *et al*., 2017). To supplement gene ontology analyses, we used STRING, a database of known protein-protein interactions, to evaluate enrichment of interacting genes from our correlation analysis (Szklarczyk *et al*., 2015). Statistical analyses were conducted in R and plots were produced using the ‘tidyverse’ package (Wickham *et al*., 2019). Phylogenies were visualised using the ape package in R (Paradis *et al*., 2003; Paradis and Schliep, 2019).

### Non-coding elements associated with convergent transition to viviparity

In order to detect regulatory elements potentially underlying the transition from oviparity to viviparity, we extracted non-coding regions from our whole genome alignment. We then partitioned the whole genome alignment by scaffold using WGAbed (https://henryjuho.github.io/WGAbed/). To infer conserved non-coding genomic regions, we estimated a neutral model by extracting and using four-fold degenerate sites across all scaffolds to transform branch lengths on the time-calibrated phylogeny obtained from Rabosky et al. (2018) using a time-reversible model (REV). Conserved non-coding regions were inferred by comparing, for each genomic region, the neutral model to a conserved model defined by scaling the neutral model by a scaling factor (‘rho’) using PhastCons with the following commands: --target-coverage 0.25 --expected-length 12 --rho 0.4 (Siepel *et al*., 2005). Finally, we used phyloP to identify conserved genomic regions that may have experienced acceleration in sequence divergence in branches where transitions to viviparity are inferred to have occurred (Siepel *et al*., 2005; Pollard *et al*., 2010; Hubisz, Pollard and Siepel, 2011)(Figure 1). To do this, for every predicted conserved non-coding region, we compared the neutral model estimated from four-fold degenerate sites to a model where we specified acceleration at two foreground branches (Figure 1) using a likelihood-ratio test and the following parameters: -- msa-format MAF --method LRT --mode ACC. All p-values were corrected for multiple testing using the Bonferroni-Hochberg procedure.

To test whether non-coding regions showing evidence of accelerated evolution in foreground branches also showed evidence of being putatively regulatory elements, we used AME (Bailey *et al*., 2009; McLeay and Bailey, 2010) to determine whether these regions showed significant enrichment of binding motifs. Binding motifs were detected using curated vertebrate transcription factor databases and compared to randomized sequences using Fisher’s exact test (McLeay and Bailey, 2010). Since in-silico approaches cannot readily and reliably detect genes that are regulated by putative regulatory elements, we followed the approach reported in Sackton et al. (2019) and assumed that putative regulatory elements in *cis* may regulate genes closest to them. To determine genes closest to accelerated putative regulatory elements, we used the bedtools ‘closest’ function and surveyed the number of accelerated putative regulatory elements nearby each gene (Quinlan and Hall, 2010).

## Acknowledgements

LY was supported by a University of St Andrews studentship. MGR, CMG & YSL are supported by a Leverhulme research grant RPG-2020-181, by a Programa de Apoyo a Proyectos de Investigación e Inovación Tecnológica (PAPIIT) research grant (PAPIIT IN210718) and by a Consejo Nacional de Ciencia y Tecnología (CONACyT) Ciencia de Frontera research grant A1-S-33467. PT and bioinformatics and computational biology analyses were supported by the University of St Andrews Bioinformatics Unit (AMD3BIOINF), funded by Wellcome Trust ISSF award 105621/Z/14/Z. Additional HPC (Crop Diversity) were awarded as part of a BBSRC 18ALERT grant (BB/S019669/1).

## Data availability

Genomes assembled in this project, including raw reads and annotation are available from NCBI using the accession codes in Supplementary Table 1. Scripts used to generate the assemblies and annotation can be found here: https://github.com/peterthorpe5/fish_genome_assembly. Scripts used in the comparative analysis of convergent evolution can be found here: https://github.com/LeebanY/Convergent-evolution-of-viviparity-in-Cyprinodontiformes.

---

**Supplementary Figure 1:**
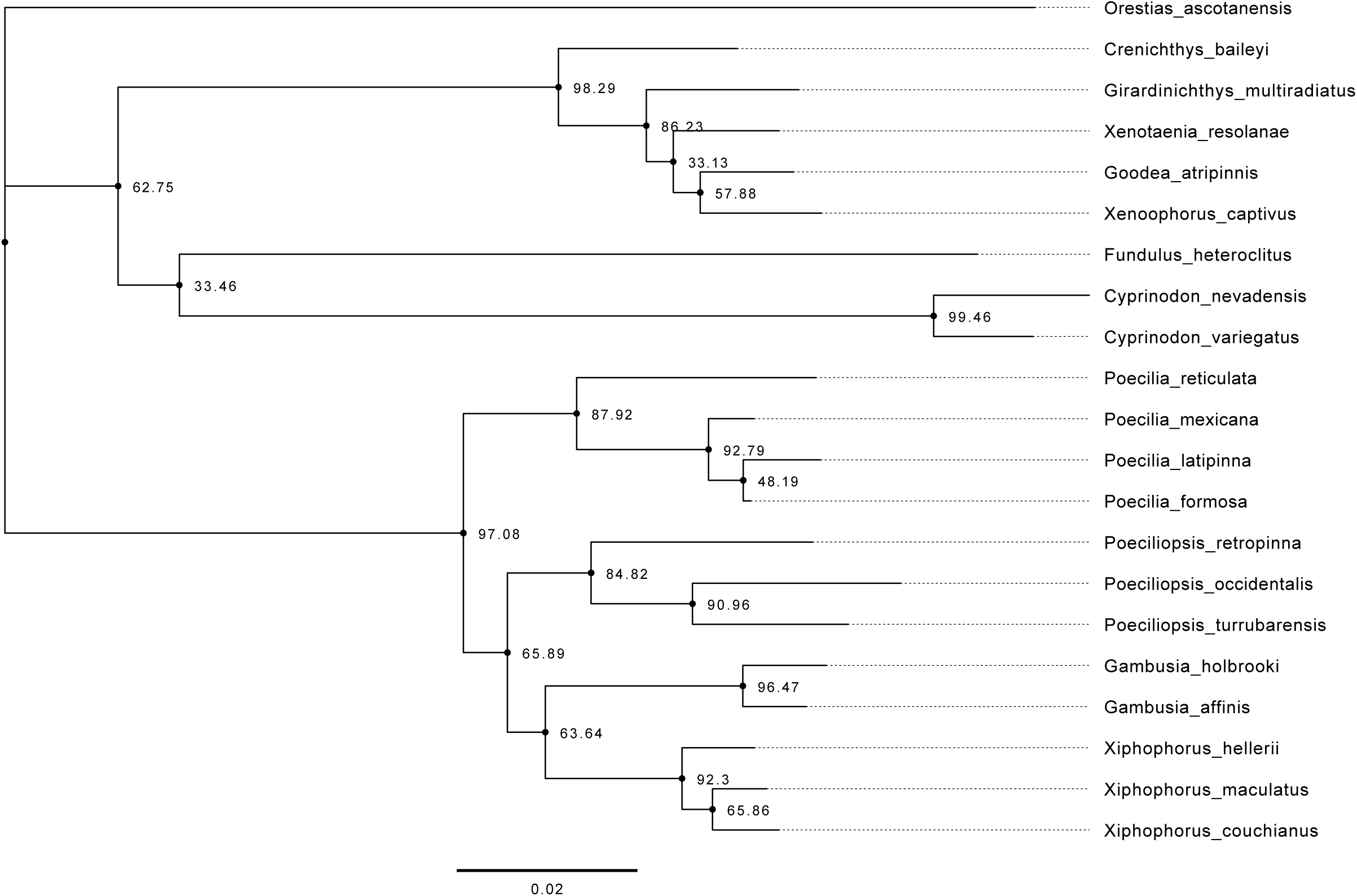
Species tree inferred using 1,044 single-copy orthologs. Gene concordance factors (shown for each node) estimated using 16,941 gene trees. Branch length represented by scale bar.

**Supplementary Figure 2:**
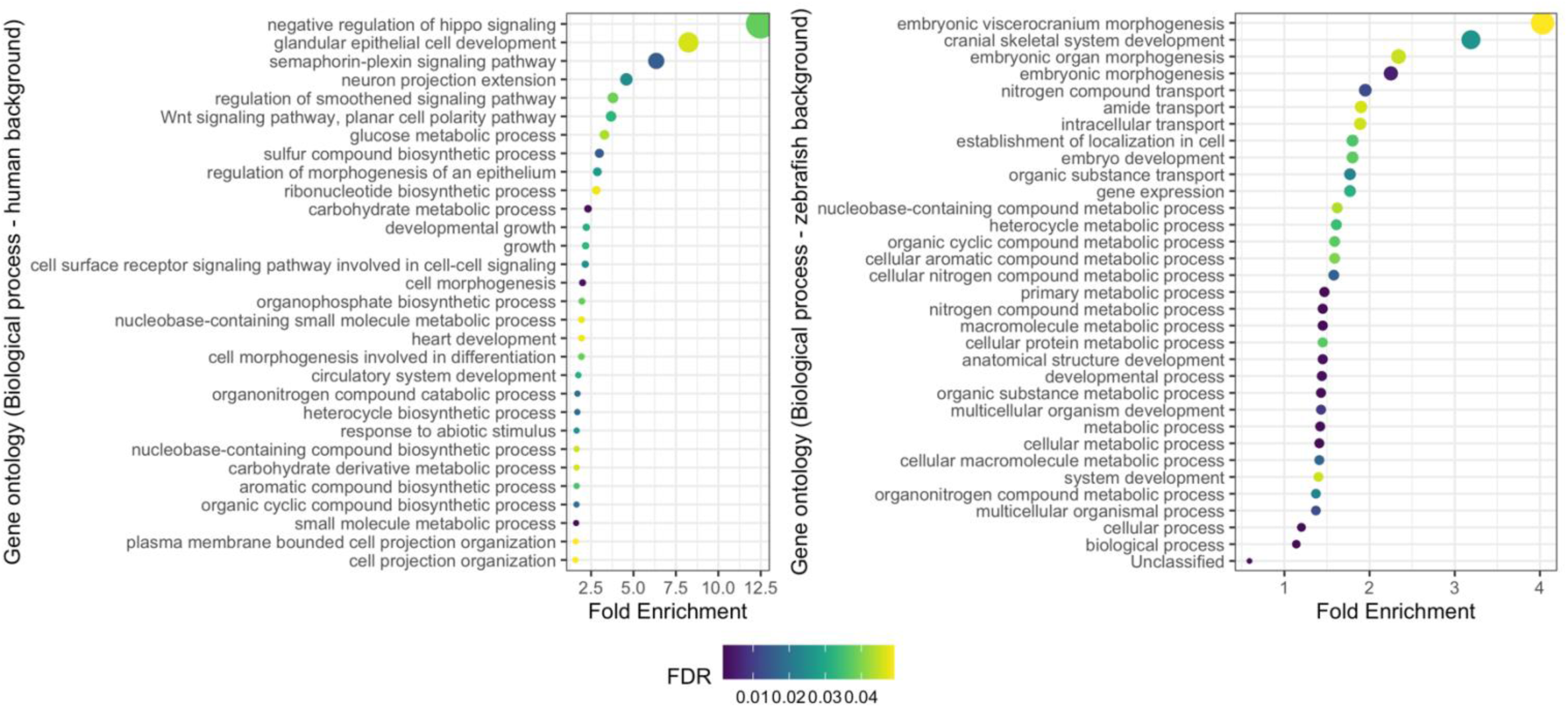
Gene ontology for biological processes of genes with a convergent amino acid change in viviparous foreground branches. Enrichment of biological processes against a human (left) and a zebrafish background/database (right) are plotted. Colours in both plots represent false discovery rates and size of points also indicate fold enrichment.

**Supplementary Figure 3:**
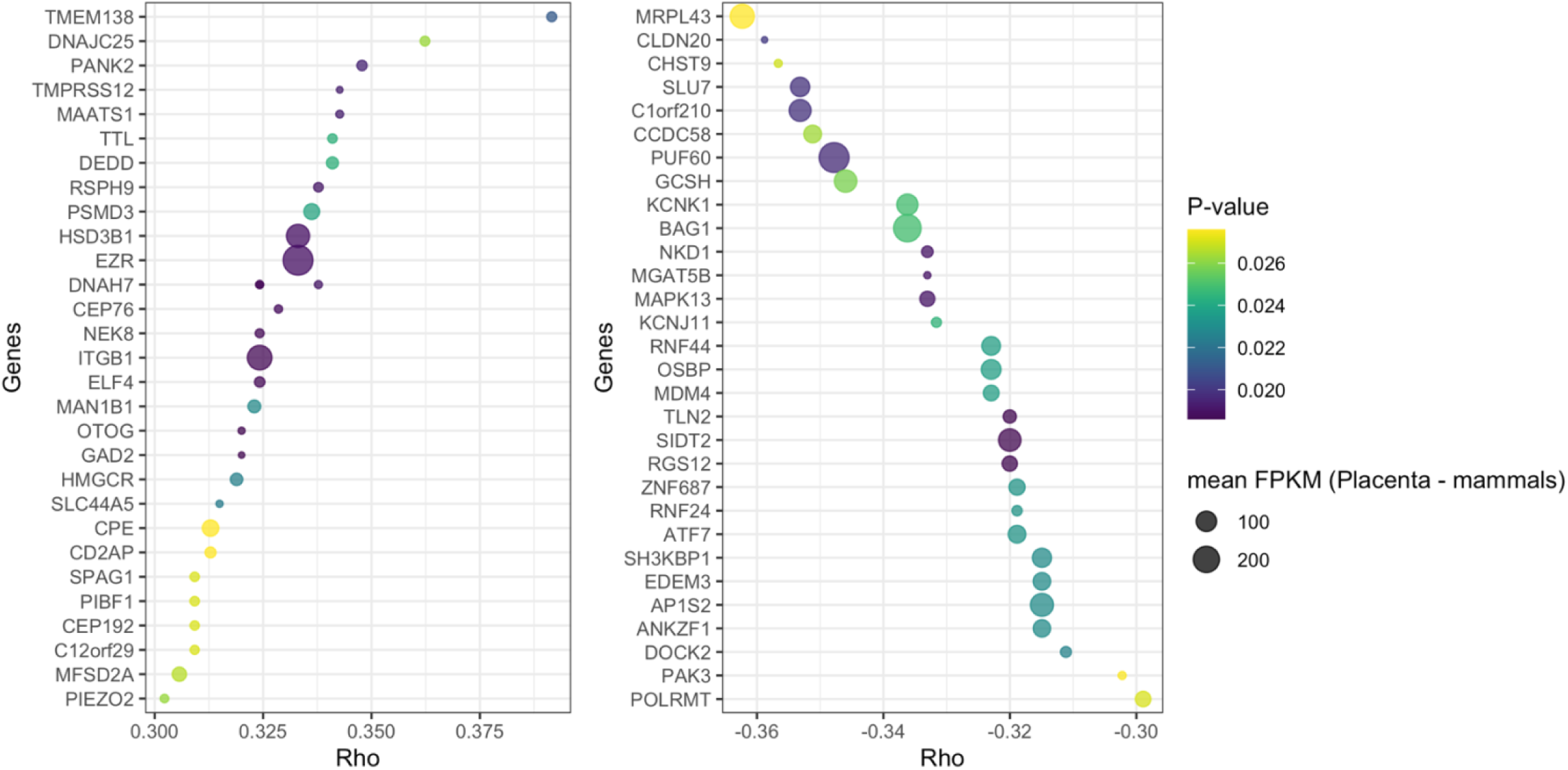
Genes undergoing convergent shifts in evolutionary rate in viviparous foreground branches. The top 30 genes with fastest rates of protein evolution (left), and the top 30 genes with slowest rates of protein evolution (right) are shown. Rho indicates the correlation between change in trait (reproductive mode) and relative rate of protein evolution. Here, points are coloured by p-value from the correlation analysis and the size of each point is indicative of mean FPKM (fragments per kilobase per million reads) in placental tissue across 14 mammal species.

**Supplementary Figure 4:**
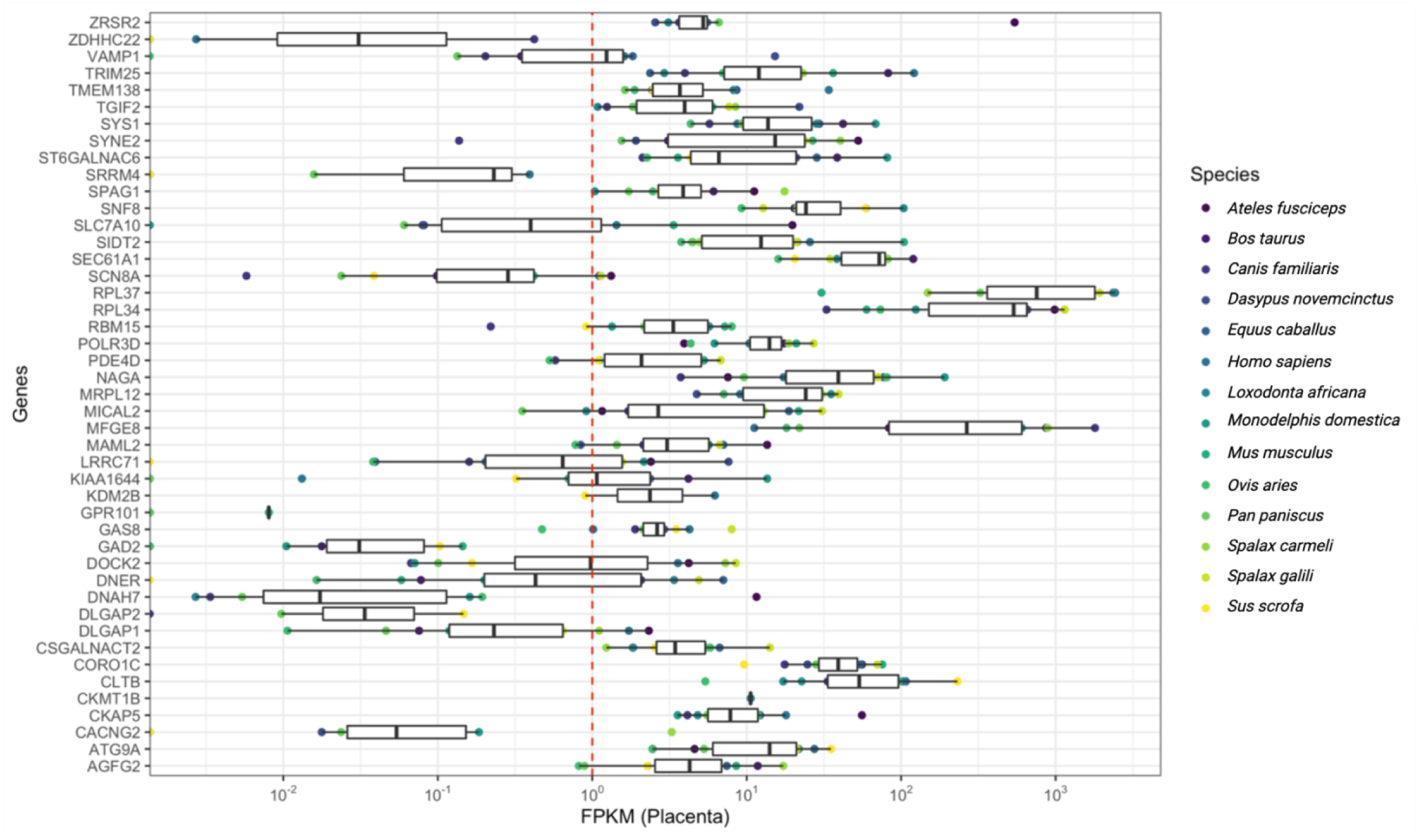
Mean placental expression in 14 mammals of genes showing signals of convergent evolution in foreground branches. Expression is represented as mean fragments per kilobase of transcript per million mapped reads (FPKM) for each gene. Coloured points represent mean FPKM for each respective gene and for a given species. Red dotted line indicates FPKM>1 cut-off. X axis is log^10^ scaled.

## References

Ali, R.H., Bogusz, M., and Whelan, S., 2019. Identifying clusters of high confidence homologies in multiple sequence alignments. Molecular biology and evolution, 36(10), pp.2340–2351.

Andrews, S. (2010) FastQC, Babraham Bioinformatics.

Anteby, E.Y., Natanson-Yaron, S., Hamani, Y., Sciaki, Y., Goldman-Wohl, D., Greenfield, C., Ariel, I., and Yagel, S., 2005. Fibroblast growth factor-10 and fibroblast growth factor receptors 1–4: expression and peptide localization in human decidua and placenta. European Journal of Obstetrics & Gynecology and Reproductive Biology, 119(1), pp.27–35.

Armstrong, D.L., McGowen, M.R., Weckle, A., Pantham, P., Caravas, J., Agnew, D., Benirschke, K., Savage-Rumbaugh, S., Nevo, E., Kim, C.J., and Wagner, G.P., 2017. The core transcriptome of mammalian placentas and the divergence of expression with placental shape. Placenta, 57, pp.71–78.

Bailey, T.L., Boden, M., Buske, F.A., Frith, M., Grant, C.E., Clementi, L., Ren, J., Li, W.W., and Noble, W.S., 2009. MEME SUITE: tools for motif discovery and searching. Nucleic acids research, 37(suppl_2), pp.W202–W208.

Benjamini, Y., and Hochberg, Y., 1995. Controlling the false discovery rate: a practical and powerful approach to multiple testing. Journal of the Royal statistical society: series B (Methodological), 57(1), pp.289–300.

Berens, A.J., Hunt, J.H., and Toth, A.L., 2015. Comparative transcriptomics of convergent evolution: different genes but conserved pathways underlie caste phenotypes across lineages of eusocial insects. Molecular biology and evolution, 32(3), pp.690–703.

Blackburn, D.G., 1992. Convergent evolution of viviparity, matrotrophy, and specializations for fetal nutrition in reptiles and other vertebrates. American Zoologist, 32(2), pp.313–321.

Blackbum, D.G., 1999. Viviparity and oviparity: evolution and reproductive strategies.

Blackburn, D.G., 2015. Evolution of viviparity in squamate reptiles: reversibility reconsidered. Journal of Experimental Zoology Part B: Molecular and Developmental Evolution, 324(6), pp.473–486.

Blanchette, M., Kent, W.J., Riemer, C., Elnitski, L., Smit, A.F., Roskin, K.M., Baertsch, R., Rosenbloom, K., Clawson, H., Green, E.D., and Haussler, D., 2004. Aligning multiple genomic sequences with the threaded blockset aligner. Genome research, 14(4), pp.708–715.

Boetzer, M., Henkel, C.V., Jansen, H.J., Butler, D., and Pirovano, W., 2011. Scaffolding pre-assembled contigs using SSPACE. Bioinformatics, 27(4), pp.578–579.

Bolger, A.M., Lohse, M., and Usadel, B., 2014. Trimmomatic: a flexible trimmer for Illumina sequence data. Bioinformatics, 30(15), pp.2114–2120.

Burkin, H.R., Rice, M., Sarathy, A., Thompson, S., Singer, C.A., and Buxton, I.L., 2013. Integrin upregulation and localization to focal adhesion sites in pregnant human myometrium. Reproductive Sciences, 20(7), pp.804–812.

Carroll, S.B., 2008. Evo-devo and an expanding evolutionary synthesis: a genetic theory of morphological evolution. Cell, 134(1), pp.25–36.

Castoe, T.A., de Koning, A.J., Kim, H.M., Gu, W., Noonan, B.P., Naylor, G., Jiang, Z.J., Parkinson, C.L., and Pollock, D.D., 2009. Evidence for an ancient adaptive episode of convergent molecular evolution. Proceedings of the National Academy of Sciences, 106(22), pp.8986–8991.

Chikina, M., Robinson, J.D., and Clark, N.L., 2016. Hundreds of genes experienced convergent shifts in selective pressure in marine mammals. Molecular biology and evolution, 33(9), pp.2182–2192.

Christin, P.A., Salamin, N., Savolainen, V., Duvall, M.R., and Besnard, G., 2007. C4 photosynthesis evolved in grasses via parallel adaptive genetic changes. Current biology, 17(14), pp.1241–1247.

Clementi, C., Tripurani, S.K., Large, M.J., Edson, M.A., Creighton, C.J., Hawkins, S.M., Kovanci, E., Kaartinen, V., Lydon, J.P., Pangas, S.A., and DeMayo, F.J., 2013. Activin-like kinase 2 functions in peri-implantation uterine signaling in mice and humans. PLoS genetics, 9(11), p.e1003863

Corbett-Detig, R.B., Russell, S.L., Nielsen, R., and Losos, J., 2020. Phenotypic convergence is not mirrored at the protein level in a lizard adaptive radiation. Molecular Biology and Evolution, 37(6), pp.1604–1614.

Davis, J.R., and Tapon, N., 2019. Hippo signalling during development. Development, 146(18), p.dev167106.

Dobin, A., Davis, C.A., Schlesinger, F., Drenkow, J., Zaleski, C., Jha, S., Batut, P., Chaisson, M., and Gingeras, T.R., 2013. STAR: ultrafast universal RNA-seq aligner. Bioinformatics, 29(1), pp.15–21

Dong, X.Y., Fu, X., Fan, S., Guo, P., Su, D., and Dong, J.T., 2012. Oestrogen causes ATBF1 protein degradation through the oestrogen-responsive E3 ubiquitin ligase EFP. Biochemical Journal, 444(3), pp.581–590.

Du, K., Pippel, M., Kneitz, S., Feron, R., da Cruz, I., Winkler, S., Wilde, B., Luna, E.G.A., Myers, G., Guiguen, Y., and Garcia, C.M., 2022. Genome biology of the darkedged splitfin, Girardinichthys multiradiatus, and the evolution of sex chromosomes and placentation. Genome Research, 32(3), pp.583–594.

Dutheil, J.Y., Gaillard, S., and Stukenbrock, E.H., 2014. MafFilter: a highly flexible and extensible multiple genome alignment files processor. BMC genomics, 15(1), pp.1–10.

Van Dyke, J.U., Brandley, M.C., and Thompson, M.B., 2014. The evolution of viviparity: molecular and genomic data from squamate reptiles advance understanding of live birth in amniotes. Reproduction, 147(1), pp.R15–R26.

Emera, D., Casola, C., Lynch, V.J., Wildman, D.E., Agnew, D., and Wagner, G.P., 2012. Convergent evolution of endometrial prolactin expression in primates, mice, and elephants through the independent recruitment of transposable elements. Molecular biology and evolution, 29(1), pp.239–247.

Emms, D.M., and Kelly, S., 2019. OrthoFinder: phylogenetic orthology inference for comparative genomics. Genome biology, 20(1), pp.1–14.

Fullerton, P.T., Monsivais, D., Kommagani, R., and Matzuk, M.M., 2017. Follistatin is critical for mouse uterine receptivity and decidualization. Proceedings of the National Academy of Sciences, 114(24), pp.E4772–E4781.

Furness, A.I., Pollux, B.J., Meredith, R.W., Springer, M.S., and Reznick, D.N., 2019. How conflict shapes evolution in poeciliid fishes. Nature communications, 10(1), pp.1–12.

Gack, M.U., Shin, Y.C., Joo, C.H., Urano, T., Liang, C., Sun, L., Takeuchi, O., Akira, S., Chen, Z., Inoue, S., and Jung, J.U., 2007. TRIM25 RING-finger E3 ubiquitin ligase is essential for RIG-I-mediated antiviral activity. Nature, 446(7138), pp.916–920.

Gao, W., Sun, Y.B., Zhou, W.W., Xiong, Z.J., Chen, L., Li, H., Fu, T.T., Xu, K., Xu, W., Ma, L., and Chen, Y.J., 2019. Genomic and transcriptomic investigations of the evolutionary transition from oviparity to viviparity. Proceedings of the National Academy of Sciences, 116(9), pp.3646–3655

Gerri, C., McCarthy, A., Alanis-Lobato, G., Demtschenko, A., Bruneau, A., Loubersac, S., Fogarty, N.M., Hampshire, D., Elder, K., Snell, P., and Christie, L., 2020. Initiation of a conserved trophectoderm program in human, cow and mouse embryos. Nature, 587(7834), pp.443–447

Grewal, S., Carver, J.G., Ridley, A.J., and Mardon, H.J., 2008. Implantation of the human embryo requires Rac1-dependent endometrial stromal cell migration. Proceedings of the National Academy of Sciences, 105(42), pp.16189–16194.

Guernsey, M.W., van Kruistum, H., Reznick, D.N., Pollux, B.J., and Baker, J.C., 2020. Molecular signatures of placentation and secretion uncovered in Poeciliopsis maternal follicles. Molecular biology and evolution, 37(9), pp.2679–2690.

Haas, B.J., Salzberg, S.L., Zhu, W., Pertea, M., Allen, J.E., Orvis, J., White, O., Buell, C.R., and Wortman, J.R., 2008. Automated eukaryotic gene structure annotation using EVidenceModeler and the Program to Assemble Spliced Alignments. Genome biology, 9(1), pp.1–22.

Haas, B.J., Papanicolaou, A., Yassour, M., Grabherr, M., Blood, P.D., Bowden, J., Couger, M.B., Eccles, D., Li, B., Lieber, M., and MacManes, M.D., 2013. De novo transcript sequence reconstruction from RNA-seq using the Trinity platform for reference generation and analysis. Nature protocols, 8(8), pp.1494–1512.

Hamada, M., Ono, Y., Asai, K., and Frith, M.C., 2017. Training alignment parameters for arbitrary sequencers with LAST-TRAIN. Bioinformatics, 33(6), pp.926–928.

Helmstetter, A.J., Papadopulos, A.S., Igea, J., Van Dooren, T.J., Leroi, A.M., and Savolainen, V., 2016. Viviparity stimulates diversification in an order of fish. Nature communications, 7(1), pp.1–7.

Hibbins, M.S., Gibson, M.J., and Hahn, M.W., 2020. Determining the probability of hemiplasy in the presence of incomplete lineage sorting and introgression. Elife, 9, p.e63753.

Home, P., Kumar, R.P., Ganguly, A., Saha, B., Milano-Foster, J., Bhattacharya, B., Ray, S., Gunewardena, S., Paul, A., Camper, S.A., and Fields, P.E., 2017. Genetic redundancy of GATA factors in the extraembryonic trophoblast lineage ensures the progression of preimplantation and postimplantation mammalian development. Development, 144(5), pp.876–888.

Hubisz, M.J., Pollard, K.S., and Siepel, A., 2011. PHAST and RPHAST: phylogenetic analysis with space/time models. Briefings in bioinformatics, 12(1), pp.41–51.

Hudson, R.R., 2002. Generating samples under a Wright–Fisher neutral model of genetic variation. Bioinformatics, 18(2), pp.337–338.

Huerta-Cepas, J., Forslund, K., Coelho, L.P., Szklarczyk, D., Jensen, L.J., Von Mering, C., and Bork, P., 2017. Fast genome-wide functional annotation through orthology assignment by eggNOG-mapper. Molecular biology and evolution, 34(8), pp.2115–2122.

Huerta-Cepas, J., Szklarczyk, D., Heller, D., Hernández-Plaza, A., Forslund, S.K., Cook, H., Mende, D.R., Letunic, I., Rattei, T., Jensen, L.J., and von Mering, C., 2019. eggNOG 5.0: a hierarchical, functionally and phylogenetically annotated orthology resource based on 5090 organisms and 2502 viruses. Nucleic acids research, 47(D1), pp.D309–D314.

Johanson, U., West, J., Lister, C., Michaels, S., Amasino, R., and Dean, C., 2000. Molecular analysis of FRIGIDA, a major determinant of natural variation in Arabidopsis flowering time. Science, 290(5490), pp.344–347.

Katoh, K., Misawa, K., Kuma, K.I., and Miyata, T., 2002. MAFFT: a novel method for rapid multiple sequence alignment based on fast Fourier transform. Nucleic acids research, 30(14), pp.3059–3066.

Katoh, K., Kuma, K.I., Toh, H., and Miyata, T., 2005. MAFFT version 5: improvement in accuracy of multiple sequence alignment. Nucleic acids research, 33(2), pp.511–518.

Katoh, K., and Standley, D.M., 2013. MAFFT multiple sequence alignment software version 7: improvements in performance and usability. Molecular biology and evolution, 30(4), pp.772–780.

Kiełbasa, S.M., Wan, R., Sato, K., Horton, P., and Frith, M.C., 2011. Adaptive seeds tame genomic sequence comparison. Genome research, 21(3), pp.487–493.

Kin, K., Maziarz, J., Chavan, A.R., Kamat, M., Vasudevan, S., Birt, A., Emera, D., Lynch, V.J., Ott, T.L., Pavlicev, M., and Wagner, G.P., 2016. The transcriptomic evolution of mammalian pregnancy: gene expression innovations in endometrial stromal fibroblasts. Genome biology and evolution, 8(8), pp.2459–2473.

Kowalczyk, A., Meyer, W.K., Partha, R., Mao, W., Clark, N.L., and Chikina, M., 2019. RERconverge: an R package for associating evolutionary rates with convergent traits. Bioinformatics, 35(22), pp.4815–4817.

van Kruistum, H., Nijland, R., Reznick, D.N., Groenen, M.A., Megens, H.J., and Pollux, B.J., 2021. Parallel Genomic changes drive repeated evolution of placentas in live-bearing fish. Molecular biology and evolution, 38(6), pp.2627–2638.

Kuittinen, H., Niittyvuopio, A., Rinne, P., and Savolainen, O., 2008. Natural variation in Arabidopsis lyrata vernalization requirement conferred by a FRIGIDA indel polymorphism. Molecular Biology and Evolution, 25(2), pp.319–329.

Kunisaki, Y., Nishikimi, A., Tanaka, Y., Takii, R., Noda, M., Inayoshi, A., Watanabe, K.I., Sanematsu, F., Sasazuki, T., Sasaki, T., and Fukui, Y., 2006. DOCK2 is a Rac activator that regulates motility and polarity during neutrophil chemotaxis. The Journal of cell biology, 174(5), pp.647–652.

Laetsch, D.R., and Blaxter, M.L., 2017. BlobTools: Interrogation of genome assemblies. F1000Research, 6(1287), p.1287.

Le, S.Q., and Gascuel, O., 2008. An improved general amino acid replacement matrix. Molecular biology and evolution, 25(7), pp.1307–1320.

Lee, K.Y., Jeong, J.W., Wang, J., Ma, L., Martin, J.F., Tsai, S.Y., Lydon, J.P., and DeMayo, F.J., 2007. Bmp2 is critical for the murine uterine decidual response. Molecular and cellular biology, 27(15), pp.5468–5478.

Lynch, V.J., Tanzer, A., Wang, Y., Leung, F.C., Gellersen, B., Emera, D., and Wagner, G.P., 2008. Adaptive changes in the transcription factor HoxA-11 are essential for the evolution of pregnancy in mammals. Proceedings of the National Academy of Sciences, 105(39), pp.14928–14933.

Lynch, V.J., and Wagner, G.P., 2008. Resurrecting the role of transcription factor change in developmental evolution. Evolution: International Journal of Organic Evolution, 62(9), pp.2131–2154.

Martin, A., and Orgogozo, V., 2013. The loci of repeated evolution: a catalog of genetic hotspots of phenotypic variation. Evolution, 67(5), pp.1235–1250.

McLeay, R.C., and Bailey, T.L., 2010. Motif Enrichment Analysis: a unified framework and an evaluation on ChIP data. BMC bioinformatics, 11(1), pp.1–11.

Mi, H., Huang, X., Muruganujan, A., Tang, H., Mills, C., Kang, D., and Thomas, P.D., 2017. PANTHER version 11: expanded annotation data from Gene Ontology and Reactome pathways, and data analysis tool enhancements. Nucleic acids research, 45(D1), pp.D183–D189.

Kent, W.J., Baertsch, R., Hinrichs, A., Miller, W., and Haussler, D., 2003. Evolution’s cauldron: duplication, deletion, and rearrangement in the mouse and human genomes. Proceedings of the National Academy of Sciences, 100(20), pp.11484–11489.

Natarajan, C., Projecto-Garcia, J., Moriyama, H., Weber, R.E., Muñoz-Fuentes, V., Green, A.J., Kopuchian, C., Tubaro, P.L., Alza, L., Bulgarella, M., and Smith, M.M., 2015. Convergent evolution of hemoglobin function in high-altitude Andean waterfowl involves limited parallelism at the molecular sequence level. PLoS genetics, 11(12), p.e1005681.

Murrell, B., Weaver, S., Smith, M.D., Wertheim, J.O., Murrell, S., Aylward, A., Eren, K., Pollner, T., Martin, D.P., Smith, D.M., and Scheffler, K., 2015. Gene-wide identification of episodic selection. Molecular biology and evolution, 32(5), pp.1365–1371.

Murugesan, S.N., Connahs, H., Matsuoka, Y., Das Gupta, M., Tiong, G.J., Huq, M., Gowri, V., Monroe, S., Deem, K.D., Werner, T., and Tomoyasu, Y., 2022. Butterfly eyespots evolved via cooption of an ancestral gene-regulatory network that also patterns antennae, legs, and wings. Proceedings of the National Academy of Sciences, 119(8), p.e2108661119.

Nakamura, T., Yamada, K.D., Tomii, K., and Katoh, K., 2018. Parallelization of MAFFT for large-scale multiple sequence alignments. Bioinformatics, 34(14), pp.2490–2492.

Natarajan, C., Hoffmann, F.G., Weber, R.E., Fago, A., Witt, C.C., and Storz, J.F., 2016. Predictable convergence in hemoglobin function has unpredictable molecular underpinnings.

Nguyen, L.T., Schmidt, H.A., Von Haeseler, A., and Minh, B.Q., 2015. IQ-TREE: a fast and effective stochastic algorithm for estimating maximum-likelihood phylogenies. Molecular biology and evolution, 32(1), pp.268–274.

Nishimura, T., Higuchi, K., Sai, Y., Sugita, Y., Yoshida, Y., Tomi, M., Wada, M., Wakayama, T., Tamura, A., Tsukita, S., and Soga, T., 2014. Fetal growth retardation and lack of hypotaurine in ezrin knockout mice. PLoS One, 9(8), p.e105423.

Nnamani, M.C., Ganguly, S., Erkenbrack, E.M., Lynch, V.J., Mizoue, L.S., Tong, Y., Darling, H.L., Fuxreiter, M., Meiler, J., and Wagner, G.P., 2016. A derived allosteric switch underlies the evolution of conditional cooperativity between HOXA11 and FOXO1. Cell reports, 15(10), pp.2097–2108.

Ornek, T., Fadiel, A., Tan, O., Naftolin, F., and Arici, A., 2008. Regulation and activation of ezrin protein in endometriosis. Human Reproduction, 23(9), pp.2104–2112.

Paradis, E., Claude, J., and Strimmer, K., 2004. APE: analyses of phylogenetics and evolution in R language. Bioinformatics, 20(2), pp.289–290.

Paradis, E., and Schliep, K., 2019. ape 5.0: an environment for modern phylogenetics and evolutionary analyses in R. Bioinformatics, 35(3), pp.526–528.

Parker, J., Tsagkogeorga, G., Cotton, J.A., Liu, Y., Provero, P., Stupka, E., and Rossiter, S.J., 2013. Genome-wide signatures of convergent evolution in echolocating mammals. Nature, 502(7470), pp.228–231.

Peng, J., Fullerton Jr, P.T., Monsivais, D., Clementi, C., Su, G.H., and Matzuk, M.M., 2015. Uterine activin-like kinase 4 regulates trophoblast development during mouse placentation. Molecular Endocrinology, 29(12), pp.1684–1693.

Peng, J., Monsivais, D., You, R., Zhong, H., Pangas, S.A., and Matzuk, M.M., 2015. Uterine activin receptor-like kinase 5 is crucial for blastocyst implantation and placental development. Proceedings of the National Academy of Sciences, 112(36), pp.E5098–E5107.

Pollard, K.S., Hubisz, M.J., Rosenbloom, K.R., and Siepel, A., 2010. Detection of nonneutral substitution rates on mammalian phylogenies. Genome research, 20(1), pp.110–121.

Pollux, B.J.A., Meredith, R.W., Springer, M.S., Garland, T., and Reznick, D.N., 2014. The evolution of the placenta drives a shift in sexual selection in livebearing fish. Nature, 513(7517), pp.233–236.

Prjibelski, A., Antipov, D., Meleshko, D., Lapidus, A., and Korobeynikov, A., 2020. Using SPAdes de novo assembler. Current protocols in bioinformatics, 70(1), p.e102.

Quinlan, A.R., and Hall, I.M., 2010. BEDTools: a flexible suite of utilities for comparing genomic features. Bioinformatics, 26(6), pp.841–842.

Rabosky, D.L., Chang, J., Cowman, P.F., Sallan, L., Friedman, M., Kaschner, K., Garilao, C., Near, T.J., Coll, M., and Alfaro, M.E., 2018. An inverse latitudinal gradient in speciation rate for marine fishes. Nature, 559(7714), pp.392–395.

Rambaut, A., and Grass, N.C., 1997. Seq-Gen: an application for the Monte Carlo simulation of DNA sequence evolution along phylogenetic trees. Bioinformatics, 13(3), pp.235–238.

Ranwez, V., Douzery, E.J., Cambon, C., Chantret, N., and Delsuc, F., 2018. MACSE v2: toolkit for the alignment of coding sequences accounting for frameshifts and stop codons. Molecular biology and evolution, 35(10), pp.2582–2584.

Recknagel, H., Carruthers, M., Yurchenko, A.A., Nokhbatolfoghahai, M., Kamenos, N.A., Bain, M.M., and Elmer, K.R., 2021. The functional genetic architecture of egg-laying and live-bearing reproduction in common lizards. Nature Ecology & Evolution, 5(11), pp.1546–1556.

Recknagel, H., and Elmer, K.R., 2019. Differential reproductive investment in co-occurring oviparous and viviparous common lizards (Zootoca vivipara) and implications for life-history trade-offs with viviparity. Oecologia, 190(1), pp.85–98.

Recknagel, H., Kamenos, N.A., and Elmer, K.R., 2018. Common lizards break Dollo’s law of irreversibility: genome-wide phylogenomics support a single origin of viviparity and re-evolution of oviparity. Molecular Phylogenetics and Evolution, 127, pp.579–588.

Ritchie, M.G., Webb, S.A., Graves, J.A., Magurran, A.E., and Macias Garcia, C., 2005. Patterns of speciation in endemic Mexican Goodeid fish: sexual conflict or early radiation?. Journal of Evolutionary Biology, 18(4), pp.922–929.

Roth, O., Solbakken, M.H., Tørresen, O.K., Bayer, T., Matschiner, M., Baalsrud, H.T., Hoff, S.N.K., Brieuc, M.S.O., Haase, D., Hanel, R., and Reusch, T.B., 2020. Evolution of male pregnancy associated with remodeling of canonical vertebrate immunity in seahorses and pipefishes. Proceedings of the National Academy of Sciences, 117(17), pp.9431–9439.

Sackton, T.B., Grayson, P., Cloutier, A., Hu, Z., Liu, J.S., Wheeler, N.E., Gardner, P.P., Clarke, J.A., Baker, A.J., Clamp, M., and Edwards, S.V., 2019. Convergent regulatory evolution and loss of flight in paleognathous birds. Science, 364(6435), pp.74–78.

Saldivar Lemus, Y., Vielle-Calzada, J.P., Ritchie, M.G., and Macías Garcia, C., 2017. Asymmetric paternal effect on offspring size linked to parent-of-origin expression of an insulin-like growth factor. Ecology and evolution, 7(12), pp.4465–4474.

Schliep, K.P., 2011. phangorn: phylogenetic analysis in R. Bioinformatics, 27(4), pp.592–593.

Shimodaira, M., Nakayama, T., Sato, I., Sato, N., Izawa, N., Mizutani, Y., Furuya, K., and Yamamoto, T., 2012. Estrogen synthesis genes CYP19A1, HSD3B1, and HSD3B2 in hypertensive disorders of pregnancy. Endocrine, 42(3), pp.700–707.

Siepel, A., Bejerano, G., Pedersen, J.S., Hinrichs, A.S., Hou, M., Rosenbloom, K., Clawson, H., Spieth, J., Hillier, L.W., Richards, S., and Weinstock, G.M., 2005. Evolutionarily conserved elements in vertebrate, insect, worm, and yeast genomes. Genome research, 15(8), pp.1034–1050.

Simão, F.A., Waterhouse, R.M., Ioannidis, P., Kriventseva, E.V., and Zdobnov, E.M., 2015. BUSCO: assessing genome assembly and annotation completeness with single-copy orthologs. Bioinformatics, 31(19), pp.3210–3212.

Smith-Unna, R., Boursnell, C., Patro, R., Hibberd, J.M., and Kelly, S., 2016. TransRate: reference-free quality assessment of de novo transcriptome assemblies. Genome research, 26(8), pp.1134–1144

Smith, M.D., Wertheim, J.O., Weaver, S., Murrell, B., Scheffler, K., and Kosakovsky Pond, S.L., 2015. Less is more: an adaptive branch-site random effects model for efficient detection of episodic diversifying selection. Molecular biology and evolution, 32(5), pp.1342–1353.

Smith, S.D., Pennell, M.W., Dunn, C.W., and Edwards, S.V., 2020. Phylogenetics is the new genetics (for most of biodiversity). Trends in Ecology & Evolution, 35(5), pp.415–425.

Soares, M.J., Konno, T., and Alam, S.K., 2007. The prolactin family: effectors of pregnancy-dependent adaptations. Trends in Endocrinology & Metabolism, 18(3), pp.114–121.

Stanke, M., Keller, O., Gunduz, I., Hayes, A., Waack, S., and Morgenstern, B., 2006. AUGUSTUS: ab initio prediction of alternative transcripts. Nucleic acids research, 34(suppl_2), pp.W435–W439.

Steiner, C.C., Römpler, H., Boettger, L.M., Schöneberg, T., and Hoekstra, H.E., 2009. The genetic basis of phenotypic convergence in beach mice: similar pigment patterns but different genes. Molecular Biology and Evolution, 26(1), pp.35–45.

Steinhart, Z., and Angers, S., 2018. Wnt signaling in development and tissue homeostasis. Development, 145(11), p.dev146589.

Stern, D.L., 2013. The genetic causes of convergent evolution. Nature Reviews Genetics, 14(11), pp.751–764.

Storz, J.F., 2016. Causes of molecular convergence and parallelism in protein evolution. Nature Reviews Genetics, 17(4), pp.239–250.

Sugawara, T., Terai, Y., Imai, H., Turner, G.F., Koblmüller, S., Sturmbauer, C., Shichida, Y., and Okada, N., 2005. Parallelism of amino acid changes at the RH1 affecting spectral sensitivity among deep-water cichlids from Lakes Tanganyika and Malawi. Proceedings of the National Academy of Sciences, 102(15), pp.5448–5453.

Suyama, M., Torrents, D., and Bork, P., 2006. PAL2NAL: robust conversion of protein sequence alignments into the corresponding codon alignments. Nucleic acids research, 34(suppl_2), pp.W609–W612.

Szklarczyk, D., Franceschini, A., Wyder, S., Forslund, K., Heller, D., Huerta-Cepas, J., Simonovic, M., Roth, A., Santos, A., Tsafou, K.P., and Kuhn, M., 2015. STRING v10: protein–protein interaction networks, integrated over the tree of life. Nucleic acids research, 43(D1), pp.D447–D452.

Tamuri, A.U., Dos Reis, M., Hay, A.J., and Goldstein, R.A., 2009. Identifying changes in selective constraints: host shifts in influenza. PLoS computational biology, 5(11), p.e1000564.

Thomas, G.W., and Hahn, M.W., 2015. Determining the null model for detecting adaptive convergence from genomic data: a case study using echolocating mammals. Molecular biology and evolution, 32(5), pp.1232–1236.

Thorpe, P., Escudero-Martinez, C.M., Cock, P.J., Eves-van den Akker, S., and Bos, J.I., 2018. Shared transcriptional control and disparate gain and loss of aphid parasitism genes. Genome biology and evolution, 10(10), pp.2716–2733.

Walker, B.J., Abeel, T., Shea, T., Priest, M., Abouelliel, A., Sakthikumar, S., Cuomo, C.A., Zeng, Q., Wortman, J., Young, S.K., and Earl, A.M., 2014. Pilon: an integrated tool for comprehensive microbial variant detection and genome assembly improvement. PloS one, 9(11), p.e112963.

Warren, W.C., García-Pérez, R., Xu, S., Lampert, K.P., Chalopin, D., Stöck, M., Loewe, L., Lu, Y., Kuderna, L., Minx, P., and Montague, M.J., 2018. Clonal polymorphism and high heterozygosity in the celibate genome of the Amazon molly. Nature ecology & evolution, 2(4), pp.669–679.

Webb, S.A., Graves, J.A., Macias-Garcia, C., Magurran, A.E., Foighil, D.Ó., and Ritchie, M.G., 2004. Molecular phylogeny of the livebearing Goodeidae (Cyprinodontiformes). Molecular phylogenetics and evolution, 30(3), pp.527–544.

Wertheim, J.O., Murrell, B., Smith, M.D., Kosakovsky Pond, S.L., and Scheffler, K., 2015. RELAX: detecting relaxed selection in a phylogenetic framework. Molecular biology and evolution, 32(3), pp.820–832.

Whittington, C.M., Griffith, O.W., Qi, W., Thompson, M.B., and Wilson, A.B., 2015. Seahorse brood pouch transcriptome reveals common genes associated with vertebrate pregnancy. Molecular Biology and Evolution, 32(12), pp.3114–3131.

Wickham, H., Averick, M., Bryan, J., Chang, W., McGowan, L.D.A., François, R., Grolemund, G., Hayes, A., Henry, L., Hester, J., and Kuhn, M., 2019. Welcome to the Tidyverse. Journal of open source software, 4(43), p.1686.

Wittkopp, P.J., and Kalay, G., 2012. Cis-regulatory elements: molecular mechanisms and evolutionary processes underlying divergence. Nature Reviews Genetics, 13(1), pp.59–69.

Wourms, J.P., and Lombardi, J., 1992. Reflections on the evolution of piscine viviparity. American Zoologist, 32(2), pp.276–293.

Yusuf, L., Heatley, M.C., Palmer, J.P., Barton, H.J., Cooney, C.R., and Gossmann, T.I., 2020. Noncoding regions underpin avian bill shape diversification at macroevolutionary scales. Genome research, 30(4), pp.553–565.

Zhang, J., 2003. Parallel functional changes in the digestive RNases of ruminants and colobines by divergent amino acid substitutions. Molecular biology and evolution, 20(8), pp.1310–1317

Zhang, P., Elabd, S., Hammer, S., Solozobova, V., Yan, H., Bartel, F., Inoue, S., Henrich, T., Wittbrodt, J., Loosli, F., and Davidson, G., 2015. TRIM25 has a dual function in the p53/Mdm2 circuit. Oncogene, 34(46), pp.5729–5738.

Zou, Z., and Zhang, J., 2015. No genome-wide protein sequence convergence for echolocation. Molecular Biology and Evolution, 32(5), pp.1237–1241.

